# Template-assisted covalent modification of DCAF16 underlies activity of BRD4 molecular glue degraders

**DOI:** 10.1101/2023.02.14.528208

**Authors:** Yen-Der Li, Michelle W. Ma, Muhammad Murtaza Hassan, Moritz Hunkeler, Mingxing Teng, Kedar Puvar, Ryan Lumpkin, Brittany Sandoval, Cyrus Y. Jin, Scott B. Ficarro, Michelle Y. Wang, Shawn Xu, Brian J. Groendyke, Logan H. Sigua, Isidoro Tavares, Charles Zou, Jonathan M. Tsai, Paul M. C. Park, Hojong Yoon, Felix C. Majewski, Jarrod A. Marto, Jun Qi, Radosław P. Nowak, Katherine A. Donovan, Mikołaj Słabicki, Nathanael S. Gray, Eric S. Fischer, Benjamin L. Ebert

## Abstract

Small molecules that induce protein-protein interactions to exert proximity-driven pharmacology such as targeted protein degradation are a powerful class of therapeutics^1-3^. Molecular glues are of particular interest given their favorable size and chemical properties and represent the only clinically approved degrader drugs^4-6^. The discovery and development of molecular glues for novel targets, however, remains challenging. Covalent strategies could in principle facilitate molecular glue discovery by stabilizing the neo-protein interfaces. Here, we present structural and mechanistic studies that define a *trans*-labeling covalent molecular glue mechanism, which we term “template-assisted covalent modification”. We found that a novel series of BRD4 molecular glue degraders act by recruiting the CUL4^DCAF16^ ligase to the second bromodomain of BRD4 (BRD4_BD2_). BRD4_BD2_, in complex with DCAF16, serves as a structural template to facilitate covalent modification of DCAF16, which stabilizes the BRD4-degrader-DCAF16 ternary complex formation and facilitates BRD4 degradation. A 2.2 Å cryo-electron microscopy structure of the ternary complex demonstrates that DCAF16 and BRD4_BD2_ have pre-existing structural complementarity which optimally orients the reactive moiety of the degrader for DCAF16_Cys58_ covalent modification. Systematic mutagenesis of both DCAF16 and BRD4_BD2_ revealed that the loop conformation around BRD4_His437_, rather than specific side chains, is critical for stable interaction with DCAF16 and BD2 selectivity. Together our work establishes “template-assisted covalent modification” as a mechanism for covalent molecular glues, which opens a new path to proximity driven pharmacology.

## Introductions

Molecular glue degraders have emerged as a powerful therapeutic modality, as demonstrated by the clinical successes of thalidomide analogs in the treatment of hematological malignancies^4,5^. These small molecule degraders stabilize the protein-protein interface between ubiquitin ligases and disease-relevant neosubstrates, resulting in ubiquitination and proteasomal degradation of the targets^6^. Unlike traditional occupancy-driven pharmacology of inhibitors, the event-driven pharmacology of degraders can result in more potent and sustained drug activity^7^. The elimination of target proteins by molecular glue degraders decreases both enzymatic and scaffold function of target proteins, leading to differentiated pharmacology and often superior inhibition of protein function^8^. Moreover, molecular glue degraders hold the potential to target proteins that do not have ligandable pockets and are considered difficult to drug, including transcription factors^9^.

The clinical efficacy of thalidomide derived drugs, such as lenalidomide, and the broad utility of targeted protein degradation in research and drug discovery has inspired numerous efforts to explore proximity-driven pharmacology^10-12^. While bi-functional molecules such as PROTACs can lead to rapid proof of concept and highly potent chemical probes, molecular glues are favorable for clinical development due to reduced size and overall chemical properties^6,9^. Despite these advantages, to date, only a small number of ubiquitin ligases have been exploited by molecular glue degraders, including CRBN^1,2^, DCAF15^13^ and DDB1^14-16^. Other proximity-driven approaches lack molecular glues. Covalency has the potential not only to aid the discovery of molecular glues, but also to impart improved efficacy through strengthening of the interface^17^. Chemo-proteomic studies have indeed identified putative covalent molecular glues^18,19^, but it remains to be shown mechanistically whether these molecules truly act as molecular glues and whether general principles can be derived to aid future discovery.

In this study, we demonstrate that a set of derivatives of JQ1^20^, a non-degrading inhibitor of BRD4, act as molecular glue degraders. Using genetic screens, biochemical analyses, medicinal chemistry, structural studies, and systematic mutagenesis, we elucidate the mechanism of action for a novel class of degraders that act through template-assisted covalent modification of DCAF16.

## Results

### JQ1-derived compounds degrade BRD4 via DCAF16

GNE-0011 (GNE11) is a derivative of the inhibitor JQ1 that has been reported to degrade BRD4^21,22^. To characterize the activity of GNE11, we generated and optimized a fluorescent reporter assay for BRD4 stability (Extended Data Fig. 1a, b). We found that GNE11 induces selective degradation of the second bromodomain of BRD4 (BRD4_BD2_) with a maximal depth of degradation at 16 hours (D_max /16 h_) of ~50% (Extended Data Fig. 1c), indicating that the BRD4_BD2_ domain, but not the first bromodomain of BRD4 (BRD4_BD1_) domain, serves as the degron for drug-mediated degradation. Through synthesis of a series of GNE11 structural analogs, we discovered an acrolein analog, TMX1, that exhibited more potent degradation of BRD4 (Fig. 1a, b, and Extended Data Fig. 1d), while maintaining selectivity for BRD4_BD2_ (D_max/16h_ ~80%) (Extended Data Fig. 1e). To examine the specificity of TMX1, we performed quantitative proteome-wide mass spectrometry in K562 cells after treatment with TMX1 for 5 hours. BRD4 was the primary degradation target with a more minor effect on two of the other BET family proteins, BRD2 and BRD3 (Fig. 1c). Treatment with JQ1, which lacks the acrolein moiety of TMX1, did not alter the abundance of BRD2, BRD3, or BRD4 (Extended Data Fig. 1f). In accordance with a ubiquitin mediated mechanism, BRD4 degradation induced by either TMX1 or GNE11, was rescued by inhibition of the proteasome with MG132, inhibition of the ubiquitin-activating enzyme UBA1 with MLN7243, or inhibition of cullin neddylation with MLN4924 (Extended Data Fig. 1g).

**Figure 1.**
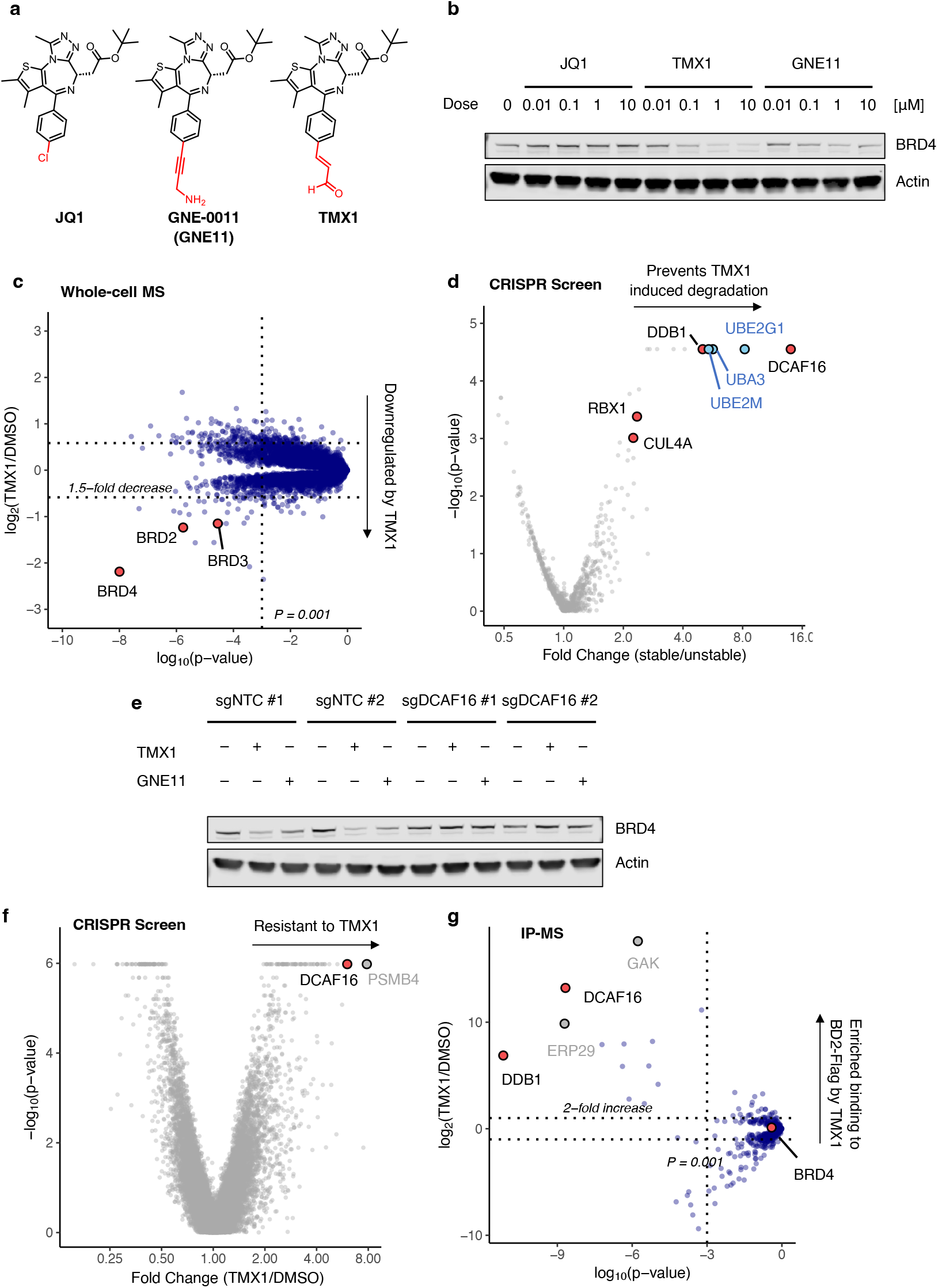
JQ1-derived compounds degrade BRD4 via DCAF16. **a**. Chemical structures of JQ1, GNE11 and TMX1. **b**. Western blots of BRD4 degradation in K562 cells treated with DMSO or different concentrations of JQ1, TMX1 and GNE11 for 16 h. **c**. Quantitative whole proteome analysis of K562 cells after treatment with TMX1 at 0.5 μM (n=2) or DMSO (n=3) for 5 h. **d**. Ubiquitin Proteasome System (UPS)-focused CRISPR degradation screen for BRD4_BD2_- eGFP stability in K562-Cas9 cells treated with TMX1 at 1 μM for 16 h (n=2). **e**. Western blots of BRD4 degradation in DCAF16 and non-targeting control (NTC) sgRNA infected K562-Cas9 cells treated with DMSO, TMX1 at 1 μM, GNE11 at 1 μM for 16 h. **f**. Genome-wide CRISPR resistance screen in K562-Cas9 cells after treatment with TMX1 at 0.1 μM (n=3) or DMSO (n=3) for 14 days. **g**. Flag immunoprecipitation (IP) followed by mass spectrometry in 293T cells overexpressing BRD4_BD2_-Flag of cells treated with either MLN4924 plus TMX1 both at 1 μM (n=4), or MLN4924 at 1 μM only (n=4). Fold enrichment and p-values were calculated by comparing TMX1/MLN4924 treated samples to MLN4924 only control samples.

To identify the molecular machinery required for TMX1 and GNE11 mediated BRD4 degradation, we performed CRISPR-Cas9 reporter degradation screens. K562 or 293T cells expressing Cas9 and the BRD4_BD2_ reporter were transduced with a single-guide RNA (sgRNA) library targeting genes in the ubiquitin-proteasome-system^14^ and then sorted for cells with increased and decreased levels of BRD4_BD2_-eGFP after drug treatment (Extended Data Fig. 2a). The screen revealed that TMX1 or GNE11 induced reporter degradation requires DCAF16, DDB1, RBX1 and CUL4A (Fig. 1d, Extended Data Fig. 2b-d). In engineered K562 cells with complete genetic knockout of DCAF16, treatment with TMX1 or GNE11 did not cause BRD4 degradation (Fig. 1e). To corroborate these findings, we performed a CRISPR-Cas9 resistance screen to identify genes required for TMX1- and GNE11-induced cellular toxicity (Extended Data Fig. 3a). sgRNAs against DCAF16 were again the most enriched, suggesting that loss of DCAF16 caused TMX1 or GNE11 resistance (Fig. 1f, and Extended Data Fig. 3b), while DCAF16 was not required for JQ1-induced cellular toxicity (Extended Data Fig. 3c). We validated that sgRNAs targeting DCAF16 confer resistance to the degraders in a competitive growth assay (Extended Data Fig. 3d). Consistent with cellular and genetic data, immunoprecipitation mass spectrometry (IP-MS) experiments with BRD4_BD2_ as a bait confirm a direct and specific compound dependent interaction with DCAF16 (Fig. 1g, and Extended Data Fig. 4a). Collectively, these data indicate that TMX1 and GNE11 act through RBX1-CUL4-DDB1-DCAF16 (CRL4^DCAF16^) ubiquitin ligase dependent degradation of BRD4.

### Covalent recruitment of DCAF16 to BRD4_BD2_

To determine the mechanism of DCAF16 recruitment, we sought to reconstitute the BRD4-DCAF16 interaction in a fully recombinant system. We developed a time-resolved fluorescence energy transfer (TR-FRET) assay (Extended Data Fig. 4b) and observed a tighter TMX1-induced interaction between DDB1-DCAF16 and BRD4_BD2_ compared to BRD4_BD1_, supporting the finding that the BD2 domain is the primary degron for TMX1-mediated degradation (Fig. 2a). We repeated a similar TR-FRET experiment with GNE11 and observed similar trends, but found that the BRD4-DCAF16 interaction was much weaker compared to TMX1 (Extended Data Fig. 4c), consistent with the lower potency of GNE11 as a BRD4 degrader. These findings suggest that TMX1 functions as a molecular glue to recruit DCAF16 selectively to BRD4_BD2_, causing degradation of BRD4.

**Figure 2.**
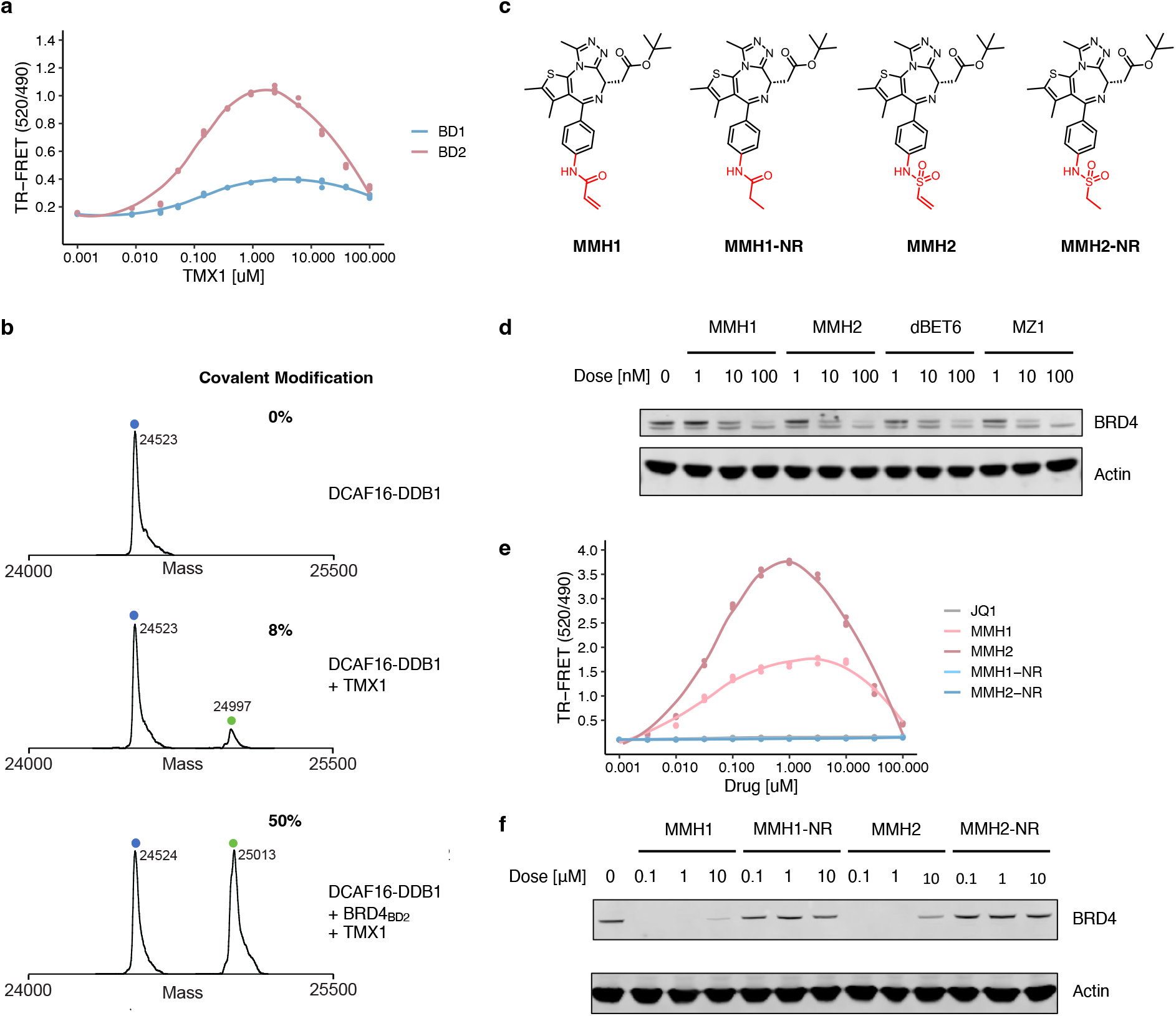
Covalent recruitment of DCAF16 to BRD4_BD2_ and optimized electrophilic warheads. **a**. TR-FRET signal for DDB1-DCAF16-BODIPY to BRD4_BD1_-terbium or BRD4_BD2_- terbium with increasing concentrations of TMX1 (n=3). **b**. Intact protein mass spectra of DDB1-DCAF16 alone, DDB1-DCAF16 co-incubated with TMX1 at 4°C for 16 h, or DDB1-DCAF16 co-incubated with TMX1 and BRD4_BD2_ at 4°C for 16 h. **c**. Chemical structures of MMH1, MMH2, MMH1-NR, MMH2-NR. **d**. Western blot of BRD4 degradation in K562 cells that were treated with DMSO or different concentrations of MMH1, MMH2, dBET6, or MZ1 for 6 h. **e**. TR-FRET signal for DDB1-DCAF16-BODIPY to BRD4_BD2_-terbium with increasing concentrations of JQ1, MMH1, MMH2, MMH1-NR and MMH2-NR (n=3). **f**. Western blots of BRD4 degradation in K562 cells that were treated with DMSO or different concentrations of MMH1, MMH1-NR, MMH2, or MMH2-NR for 16 h.

In TR-FRET experiments, the interaction of DCAF16 and BRD4_BD2_ unexpectedly decreased when the concentration of TMX1 exceeded 5μM (Fig. 2a). We also observed decreases in reporter degradation at similar compound concentrations (Extended Data Fig. 1e). This pattern, referred to as the hook effect, is commonly seen with heterobifunctional degraders in which both compound-substrate and compound-ligase interactions become saturated at high ligand concentrations^23^. Hook effects are not observed with canonical molecular glues. Since TMX1 contains an electrophilic acrolein moiety, we hypothesized that TMX1 might form a covalent bond with DCAF16, thereby providing an alternative explanation for the hook effect.

To test whether TMX1 forms a covalent bond, we incubated recombinant DCAF16-DDB1 with TMX1 and performed intact mass spectrometry. We observed minimal (8%) modification of DCAF16 (Fig. 2b). Next, to see if the ternary complex might facilitate covalent bond formation, we incubated both recombinant BRD4_BD2_ and DCAF16-DDB1 with TMX1 and performed intact mass spectrometry. With both ubiquitin ligase and substrate present in the reaction, we observed 50% modification of DCAF16 (Fig. 2b). These data suggest that TMX1 has negligible reactivity with DCAF16 alone and that the presence of BRD4_BD2_ facilitates covalent modification, perhaps because it orients the acrolein warhead for attack by the cysteine in a mechanism that we refer to as “template-assisted covalent modification.” We observe similar modification with GNE11 albeit much weaker with only a 7% DCAF16 mass shift in the presence of BRD4_BD2_ even at extended timepoints (Extended Data Fig. 4d). The weaker reactivity of GNE11 is consistent with the propargylamine, while previously shown to be reactive^24^, being a weaker electrophile. As a control, we investigated the previously reported covalent DCAF16-dependent BRD4 heterobifunctional degrader KB02-JQ1^25^, which exhibited the expected covalent modification of DCAF16 regardless of whether BRD4_BD2_ was included in the reaction (Extended Data Fig. 4e). These studies demonstrate that the JQ1-derived molecular glue degraders act through a template-assisted covalent mechanism that is distinct from heterobifunctional degraders or traditional molecular glue degraders.

### Optimized electrophilic warheads increases potency of degraders

The observation that a more reactive molecule, TMX1, demonstrated higher degradation potency than GNE11 suggests that optimization of the covalent warhead might improve the degradation activity of DCAF16-based BRD4 degraders. To test this hypothesis and to facilitate structural studies, we expanded the electrophilic chemotypes on the phenyl exit vector and characterized their BRD4_BD2_ degradation and DCAF16 recruitment activity using degradation and TR-FRET assays. We discovered an acrylamide analog, MMH1, and a vinyl sulfonamide analog, MMH2 (Fig. 2c), that both showed improved BRD4_BD2_ degradation activity (D_max, 16 h_ ~95%; and half-maximal degradation concentration at 16 hours (DC_50, 16 h_) ~1 nM) (Extended Data Fig. 5a-c) and significantly stronger DCAF16 binding as compared to TMX1 (Extended Data Fig. 5d). When comparing MMH1 and MMH2 induced degradation of BRD4 with non-covalent BRD4 heterobifunctional degraders, dBET6 and MZ1^26,27^, we found that MMH1 and MMH2 exhibit comparable BRD4 degradation (Fig. 2d, and Extended Data Fig. 5e) with more sustained activity after washout due to the covalent mechanism (Extended Data Fig. 5f).

To further confirm that covalent reactivity is critical for DCAF16 recruitment, we developed MMH1-NR and MMH2-NR, containing a non-reactive (ethyl) group and a saturated vinyl moiety, respectively (Fig. 2c). Compared to their reactive analogs, both non-reactive molecules demonstrated negligible DCAF16 recruitment (Fig. 2e) or degradation activity (Fig. 2f), indicating that covalency is required for the activity of JQ1-derived DCAF16-based BRD4 degrader. To ensure that MMH1 and MMH2 conserve the mechanism of action of TMX1 and GNE11, we repeated the BD1/BD2 degradation assay and whole-cell proteomics experiments and observed similar results (Extended Data Fig. 6a-d). Furthermore, we performed DCAF16 intact mass spectrometry experiments on MMH1 and MMH2 and found similar template-assisted covalent modifications (Extended Data Fig. 6e, f). However, the more reactive molecules, MMH1 and MMH2, also caused increased baseline, non-templated-assisted covalent labeling of DCAF16 (Extended Data Fig. 6e, f).

### BRD4_BD2_ orients MMH2 for DCAF16 modification

To understand how BRD4_BD2_ facilitates covalent modification of DCAF16, we sought to structurally characterize the ternary complex by cryo-electron microscopy (cryo-EM). Recombinant DDB1**Δ**B-DDA1-DCAF16 complex was mixed with recombinant BRD4_BD2_ and MMH2 and purified over size exclusion chromatography. A dataset was collected on a Titan Krios microscope after several rounds of grid optimization leading to a condition containing 0.011% Lauryl Maltose Neopentyl Glycol (LMNG) detergent on UltrAuFoil grids that mitigated preferred orientations of the particles (see **Methods** for details). Following several rounds of classification, a final reconstruction was refined to 2.2 Å and used for model building (Fig. 3a, b and Extended Data Fig. 7a-f, Extended Data Table 1). DDB1**Δ**B and BRD4_BD2_ were readily placed into density using high resolution structures PDB: 6Q0R and 6VIX as models. The density filling the gap between DDB1 and BRD4_BD2_ was identified as DCAF16 and a model was manually built (Fig. 3b, and Extended Data Fig. 7a-f, 8a-c, Extended Data Table 1).

**Figure 3.**
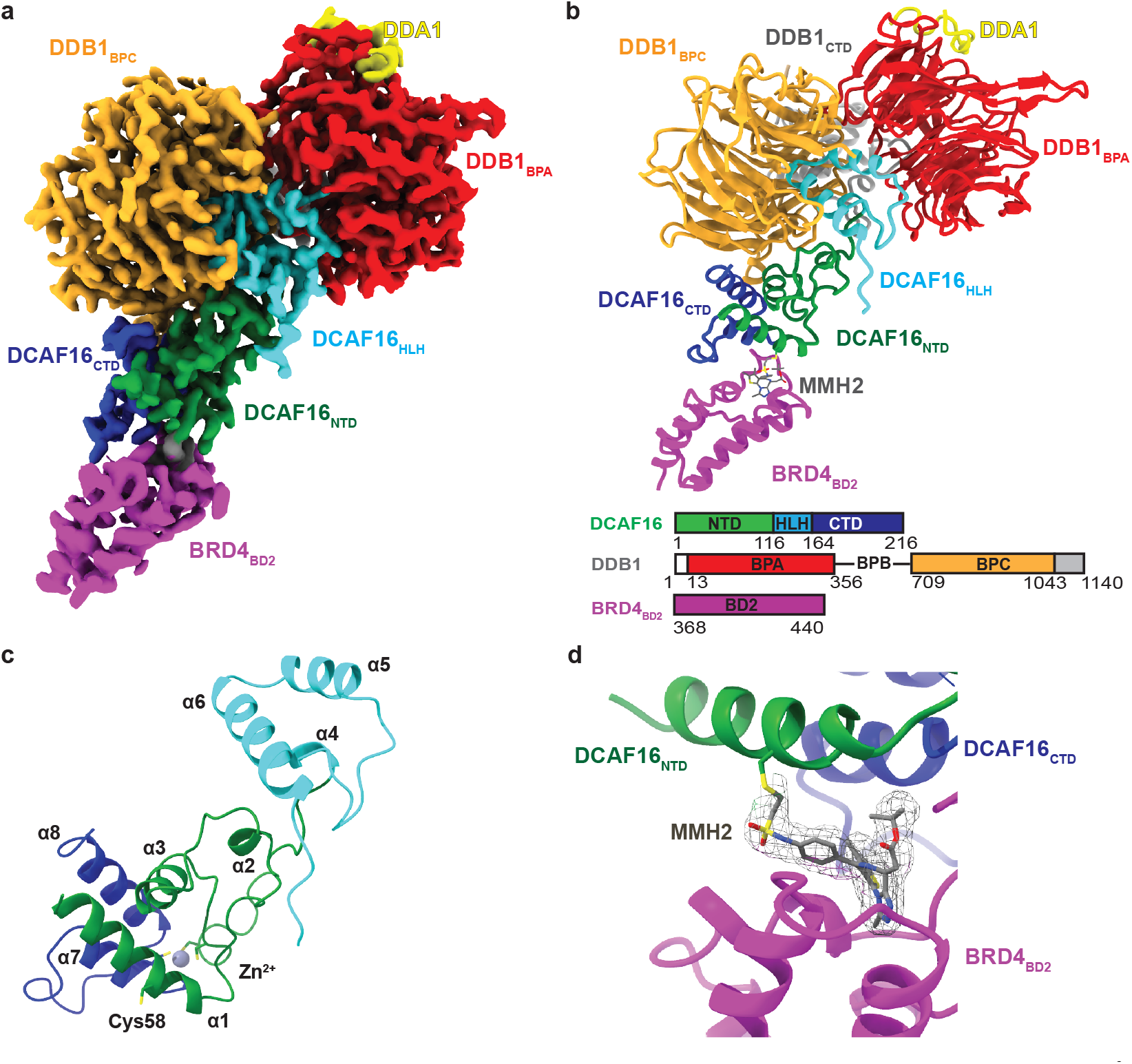
BRD4_BD2_ orients MMH2 for DCAF16 modification. **a**. 2.2 Å cryo-EM map of the DDB1**Δ**B-DDA1-DCAF16-BRD4_BD2_-MMH2 colored to indicate DDB1BPA (red), DDB1BPC (orange), DDB1CTD (gray), DDA1 (yellow), DCAF16CTD (blue), DCAF16NTD (green), DCAF16HLH (cyan), and BRD4_BD2_ (magenta). Map shown has been processed with DeepEMhancer^40^ **b**. Cartoon representation of the DDB1-DCAF16 ligase complex bound to BRD4_BD2_ and MMH2 with same coloring as the cryo-EM map. A sequence scheme for all complex partners is shown at the bottom. **c**. Cartoon representation of DCAF16 indicating secondary structure elements. **d**. Close-up of MMH2 covalently modifying DCAF16_Cys58_ with cryo-EM density around MMH2 shown as mesh.

DCAF16 folds into a structure without any homologies across the PDB or AlphaFold 2 databases^28,29^. DCAF16 is anchored to DDB1 with a helix-loop-helix (HLH) motif distinct from canonical DCAFs that occupies a similar spatial location to the DDB1-binding motif of CRBN, which is located centrally (aa 113-155). The amino-terminal and carboxy-terminal regions of DCAF16 fold into a four-helix bundle stabilized by a zinc atom forming the primary interface with BRD4_BD2_ (Fig. 3c). The first helix (α1) is followed by an extended loop towards DDB1 with a short helix (α2) packing against the HLH motif. The next helix (α3) packs on top of α1 and together with α7 and α8 forms the core of the structure. Following another extended loop and short helix (α4) back towards DDB1, the HLH-motif is formed by α5, α6 and several smaller loops filling the DDB1 cavity. Returning from the HLH-motif, another extended loop leads back to the BRD4_BD2_ interacting region forming α7, followed by a loop embracing BRD4_BD2_, and α8, completing the core structure.

DCAF16 embraces BRD4_BD2_ with major contacts contributed by α1, α7 and the loop between α7 and α8 (Fig. 3b-d, and Extended Data Fig. 8d) for a total interface area of 560 Å^2^ as assessed using the PISA server^30^. At the interface between DCAF16 and BRD4_BD2_, we observed a density representing MMH2, overlapping with the JQ1 binding site of BRD4_BD2_ (Fig. 3d, Extended Data Fig. 8d). In line with a covalent mechanism, continuous density is observed between MMH2 and Cys58 on DCAF16 (Fig. 3d), with the right geometry and distances for a covalent bond.

Additionally, key contacts between MMH2 and DCAF16 (Leu59, Lys61, Tyr62, Trp181) and BRD4_BD2_ (including Trp374, Val380, Leu385, Leu387, Tyr432, Asn433, His437), respectively, contribute to the DCAF16-BRD4_BD2_ interface (Extended Data Fig. 8e). Together, the structure and biochemical characterization support a model in which MMH2 binds BRD4_BD2_, leading to recruitment of DCAF16 and orientation of MMH2 for modification of DCAF16_Cys58_. Our data further suggest that this covalent modification of DCAF16 is necessary to stabilize the ternary complex sufficiently for ubiquitylation and consequent degradation to occur.

### DCAF16_Cys58_ is targeted by molecular glue degraders

To further corroborate the structural findings in an unbiased fashion, we performed a systematic alanine scan on all residues of DCAF16 and evaluated drug-induced BRD4_BD2_ reporter degradation in a pooled screening format (Extended Data Fig. 9a). A53R, C177A, C179A mutants scored as the top hits in those screens with all the molecular glue degraders (Fig. 4a, and Extended Data Fig. 9b-e). We validated that these mutations prevent both drug-induced BRD4 degradation and DCAF16-BRD4_BD2_ binding (Fig. 4b, c, and Extended Data Fig. 10a-f). These same three amino acids scored when we performed the screen with KB02-JQ1, a DCAF16-dependent BRD4 PROTAC^25^ (Extended Data Fig. 11a, b). These results indicate that Ala53, Cys177 and Cys179 are critical for the general E3 ubiquitin ligase function of DCAF16, but are not specific to template-assisted covalent interactions with the BRD4 molecular glue degraders. These residues are critical for DCAF16 structural integrity as Ala53 oriented towards the hydrophobic core, and Cys177 and Cys179 coordinate a structural zinc ion (Extended Data Fig. 11d, e).

**Figure 4.**
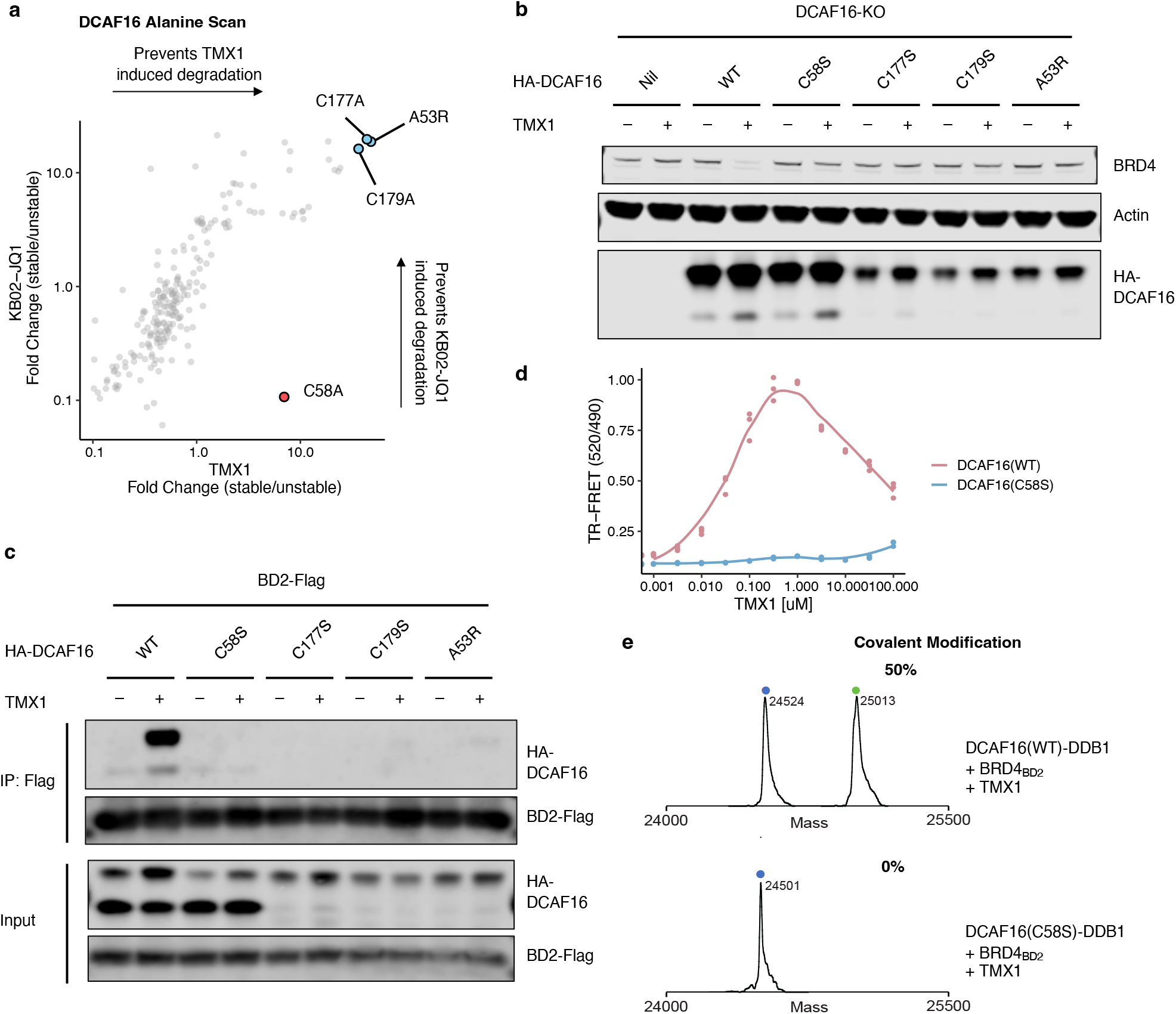
DCAF16_Cys58_ is targeted by molecular glue degraders. **a**. Correlation of fold change for two DCAF16 alanine scans in DCAF16 knockout K562 cells. The *x* axis is a degradation screen for BRD4_BD2_-eGFP upon treatment with TMX1 at 1 μM for 16 h (n=3), and the *y* axis is another degradation screen for BRD4_BD2_-eGFP upon treatment with KB02-JQ1 at 10 μM for 16 h (n=3). **b**. Western blots of BRD4 degradation in DCAF16 knockout K562 cells that were transduced with indicated HA-DCAF16 mutants, and treated with DMSO or TMX1 at 1 μM for 16 h. **c**. Flag immunoprecipitation (IP) followed by Western blots in the presence of DMSO or TMX1 at 1 μM from 293T cells transfected with indicated HA-DCAF16 mutants and BRD4_BD2_-Flag constructs. **d**. TR-FRET signal for DDB1-DCAF16(WT)- or DDB1-DCAF16(C58S)-BODIPY to BRD4_BD2_-terbium with increasing concentrations of TMX1 (n=3). **e**. Intact protein mass spectra of DDB1-DCAF16(WT) or DDB1-DCAF16(C58S) co-incubated with TMX1 and BRD4_BD2_ at 4°C for 16 h.

Only one cysteine residue, Cys58, was required exclusively for the activity of the molecular glue degraders but not for KB02-JQ1 activity (Fig. 4a, and Extended Data Fig. 9b-d, 11a). We confirmed the Cys58-selective effect on binding and degradation using co-immunoprecipitation, TR-FRET, western blots, and degradation assays (Fig. 4b-d, and Extended Data Fig. 10a-f, 11b-c). We also expressed and purified recombinant DCAF16 protein with Cys58 mutated to serine. By intact mass spectrometry the DCAF16 C58S mutant completely eliminated DCAF16-TMX1 adduct formation (Fig. 4e). We also performed intact mass spectrometry analysis on wild-type and C58S mutant DCAF16 co-incubated with BRD4_BD2_ and MMH2, showing that DCAF16 C58S mutant greatly reduces adduct formation from 95% to 20%, close to the baseline labeling efficiency of MMH2 without the presence of BRD4_BD2_ template (Extended Data Fig. 11f). Collectively, these results validate the structural insight that DCAF16_Cys58_ is the amino acid targeted for template-assisted covalent modification by the BRD4 molecular glue degraders.

### BRD4_BD2_ residues critical for interface conformation confer selectivity

Since we observed a selectivity for BRD4_BD2_, despite close homology of the BD1 and BD2 domains around the drug binding site and similar affinities for JQ1, we set out to dissect the residues on BRD4_BD2_ critical for degradation with a systematic alanine scan (Extended Data Fig. 12a). For BRD4_BD2_, His437 was the most critical residue for activity of molecular glue degraders, but not heterobifunctional degraders, indicating that it is functionally important for drug-induced DCAF16-BRD4_BD2_ recruitment (Fig. 5a, and Extended Data Fig. 12b-e, 13a-c). Known JQ1 contacting residues, including Asn433, Tyr432, Tyr390, and Trp374^20,31^, also scored as amino acids required for dBET6 and MZ1 induced degradation (Extended Data Fig. 13a, b). These findings were validated using individual alanine mutants with consistent results (Fig. 5b).

**Figure 5.**
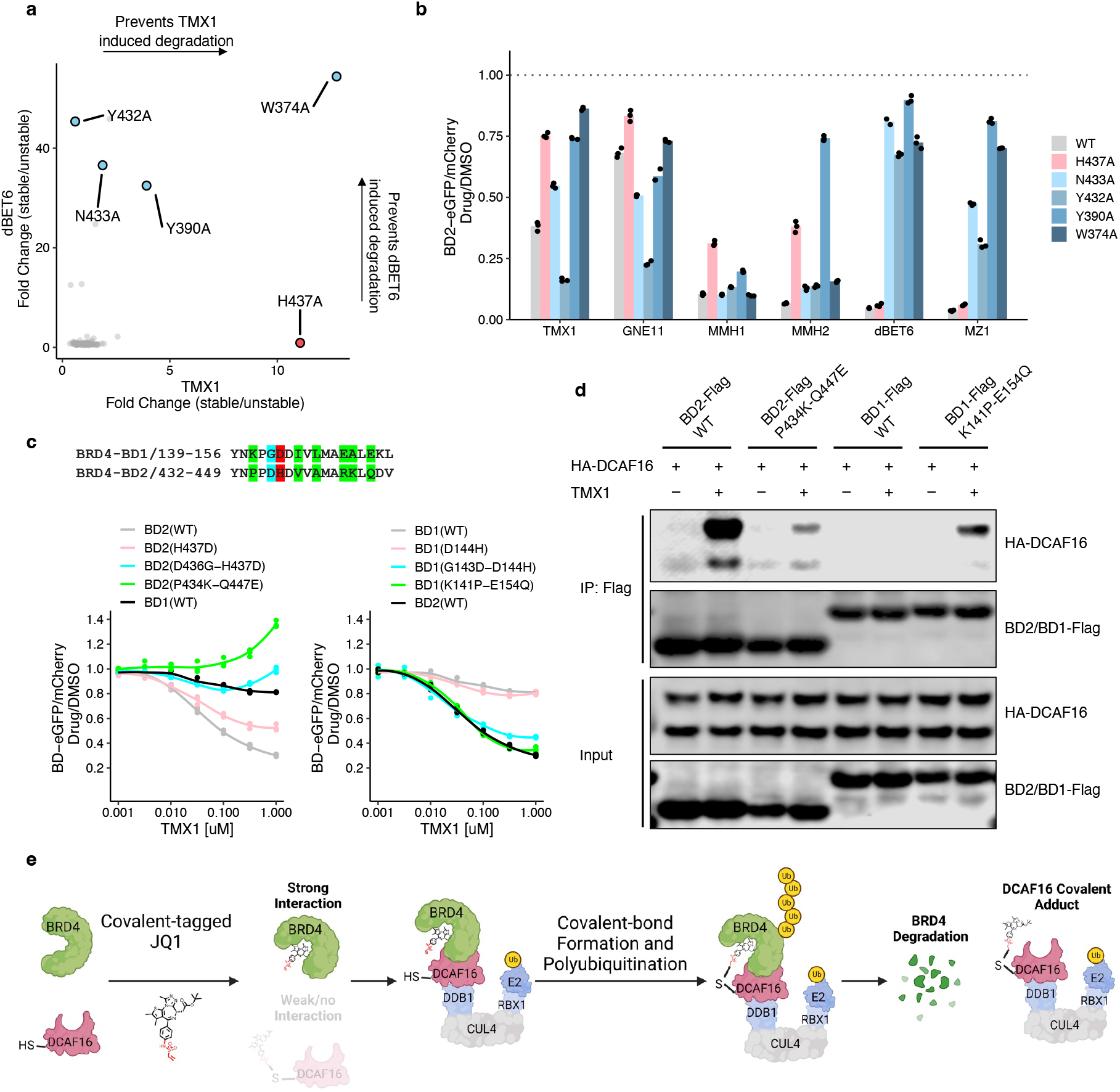
BRD4_BD2_ residues critical for interface conformation confer selectivity. **a**. Correlation of fold change for two BRD4_BD2_ alanine mutagenesis screens. The *x* axis is a degradation screen for BRD4_BD2_-eGFP in K562 cells upon treatment with TMX1 at 1 μM for 16 h (n=2), and the *y* axis is another degradation screen for BRD4_BD2_-eGFP in K562 cells upon treatment with dBET6 at 1 μM for 16 h (n=2). **b**. Flow cytometry analysis of K562 cells expressing wild-type or indicated mutant BRD4_BD2_-eGFP construct and treated with DMSO, GNE11 at 1 μM, TMX1 at 1 μM, MMH1 at 0.1 μM, MMH2 at 0.1 μM, MZ1 at 1 μM or dBET6 at 1 μM for 16 h (n=3). **c**. Flow cytometry analysis of K562 cells expressing the indicated BRD4_BD2_-eGFP, BRD4_BD1_-eGFP mutant construct and treated with increasing concentrations of TMX1 for 16 h (n=3). **d**. Flag immunoprecipitation (IP) followed by Western blots in the presence of DMSO or TMX1 at 1 μM from 293T cells transfected with HA-DCAF16 and indicated BRD4_BD2_-Flag, BRD4_BD1_-Flag mutant constructs. **e**. Schematic model for the mechanism of action of covalent BRD4 molecular glue degraders.

When comparing our structure with JQ1-bound structures of BD1 (PDB: 3MXF) or BD2 (PDB: 3ONI), the only notable differences are in the loop containing His437, closing onto the JQ1 pocket (Extended Data Fig. 13d, e). His437 contacts a carbonyl of JQ1-based degraders and potentially contributes weak interactions with Tyr62 of DCAF16 (Extended Data Fig. 8e). We therefore tested whether His437 is critical for the BD2 selectivity of covalent BRD4 molecular glue degraders. We constructed BRD4_BD1_ and BRD4_BD2_ domains, swapping the respective amino acid residues near His437. Using reporter degradation assays, we found that the BD2(D436G-H437D) and BD2(P434K-Q447E) mutants were resistant to TMX1-induced degradation, while the corresponding BD1(G143D-D144H) and BD1(K141P-E154Q) mutants gained susceptibility to TMX1-induced degradation compared with wild-type BD1 (Fig. 5c). The same amino acid swap in BD2 decreased TMX1-induced binding to DCAF16 and increased drug-induced BD1-DCAF16 binding (Fig. 5d). Given that TMX1 has comparable binding affinity to both BD1 and BD2 domains of BRD4 (Extended Data Fig. 13f), the BD2 selectivity of drug-induced degradation is likely driven by differences in protein-protein interactions between BD2 and DCAF16 and orientation of the reactive warhead with respect to DCAF16_Cys58_. While BRD4_His437_ directly contributes to binding of DCAF16, Asp436, Pro434 and Gln447 are not at the DCAF16 interface and contribute to the overall bromodomain conformation.

## Discussion

Our studies reveal a class of molecular glue degraders that act through template-assisted covalent modification and establish a mechanism for how *trans*-labeling can stabilize a molecular glue induced neo-protein protein interface, informing future discovery and design of molecular glues. Combining cellular, biochemical and structural studies, we found that the BD2 domain of BRD4, in complex with DCAF16, serves as a structural scaffold to orient the reactive moiety of a small molecule for covalent modification of DCAF16 and degradation of BRD4 (Fig. 5e). This templated reactivity has the potential to increase the affinity of other complementary protein surfaces, resulting in novel molecular glues to drive protein degradation or other biological processes. Similarly, kinase-catalyzed transfer of the electrophilic terminal phosphate group of ATP to substrate proteins can be viewed as a template (kinase)-assisted covalent modification. More broadly, it is likely that many sites observed in chemo-proteomics covalent fragment screens^32,33^, and especially those observed in unstructured regions, may be the result of a similar template-assisted mechanism in which the primary binding energy is derived from a binding partner.

The covalent property of the BRD4 degraders leads to a hook effect and more durable degradation, distinguishing covalent from non-covalent molecular glue degraders. Our studies reveal that modulating the reactivity of the electrophilic warhead can tune the activity and specificity of covalent molecular glue degraders. In the case of the JQ1-derived molecules, non-reactive molecules did not induce protein degradation, and highly reactive molecules may lack specificity. Both orientation of the covalent warhead and degree of reactivity optimizes activity and specificity. Our cryo-EM structure, with unbiased systematic mutagenesis screen and following biochemical validation, unambiguously confirmed DCAF16_Cys58_ as the amino acid targeted for covalent modification with high efficiency only when oriented by the BRD4-compound-DCAF16 ternary complex.

DCAF16 is well suited for template-assisted covalent modification since its cysteine-rich substrate binding surface is readily targeted for covalent modification, as demonstrated by our work and the prior identification of heterobifunctional degraders targeting DCAF16^25^. We determined the cryo-electron microscopy structure of the DDB1-DCAF16 ligase complex bound to BRD4_BD2_ and MMH2, providing definite proof of this molecular glue interaction. Unlike most other DCAF proteins, DCAF16 does not contain a canonical WD40 propeller^34^, and instead is a relatively unstable protein predicted to be largely unstructured. It is noteworthy that our structural studies suggest a high degree of conformational flexibility, similar to findings from studies of CRBN^35-37^, and we speculate that such structural plasticity in a ligase can facilitate glue activity.

A central challenge for the development of molecular glue degraders is the need for approaches for rational drug design and discovery^38^. In our current study, we demonstrate that the addition of electrophiles to the solvent-exposed side of JQ1, results in DCAF16-dependent covalent molecular glue degraders. The addition of electrophilic warheads to protein binders could become an effective strategy to stabilize a ternary complex and enable protein degradation when similar non-covalent molecules do not^39^. We envision that template-assisted covalent modification strategies can be exploited to facilitate future rational design and discovery of molecular glue degraders.

## Materials and Methods

### Mammalian cell culture

The human HEK293T and HEK293T-Cas9 cell lines were provided by the Genetic Perturbation Platform, Broad Institute. K562-Cas9 cell line was provided by Zuzana Tothova (Dana-Farber Cancer Institute). HEK293T and HEK293T-Cas9 cells were cultured in Dulbecco’s modified Eagle’s medium (DMEM) (Gibco), and K562-Cas9 cell lines were cultured in RPMI (Gibco), with 10% fetal bovine serum (FBS) (Invitrogen), 5% glutamine and penicillin-streptomycin (Invitrogen) at 37 °C and 5% CO2.

### Antibodies

The following antibodies were used: anti-BRD4 (Bethyl Laboratories, A301-985A100), anti-β-actin (Cell Signaling Technology, #3700), anti-Flag (Sigma-Aldrich, M8823), anti-HA (Cell Signaling Technology, #3724), anti-mouse 800CW (LI-COR Biosciences, 926-32211), anti-rabbit 680LT (LI-COR Biosciences, 925-68021).

### Compounds

JQ1 (HY-13030), dBET6 (HY-112588), MZ1 (HY-107425), KB02-JQ1 (HY-129917), and MLN4924 (HY-70062) were obtained from MedChemExpress; MLN7243 (CT-M7243) was obtained from ChemieTek; MG132 (S2619) was obtained from Selleck Chemicals.

### Plasmids

The following plasmids were used in this study: Cilantro (PGK.BsmBICloneSite.FlexibleLinker.eGFP.IRES.mCherry.cppt.EF1α.PuroR, Addgene 74450) for degradation characterization, reporter CRISPR screen, BRD4_BD2_ alanine scan and DCAF16 alanine scan; sgBFP (U6.sgRNA.cppt.SFFV.tBFP) and sgRFP (U6.sgRNA.cppt.EF1α.RFP657) for validation of DCAF16 knockout phenotypes; Mint-Flag/HA (SFFV.BsmBICloneSite.Flag/HA.cppt.EF1α.PuroR) and Ivy-Flag/HA (SFFV.Flag/HA.BsmBICloneSite.cppt.EF1α.PuroR) for co-immunoprecipitation and DCAF16 mutant transduction; pAC8-derived plasmids for protein purification; E.Coli pET100/D-TOPO for protein purification.

### Immunoblots

Cells were washed with PBS and lysed in RIPA lysis buffer (Thermo Fisher Scientific) with Halt Protease Inhibitor Cocktail (Thermo Fisher Scientific) and Benzonase (Sigma-Aldrich) for 20 min on ice. The insoluble fraction was removed by centrifugation, the protein concentration was quantified using a BCA protein assay kit (Thermo Fisher Scientific), and an equal amount of lysate was run on SDS–PAGE 4–12% Bis–Tris Protein Gels (Thermo Fisher Scientific) and then transferred to nitrocellulose membrane with a XCell II Blot Module Wet Tank Transfer System (Thermo Fisher Scientific). Membranes were blocked in Intercept (PBS) Blocking Buffer (LI-COR Biosciences) and incubated with primary antibodies overnight at 4 °C. The membranes were then washed in Tris-buffered saline with Tween-20 (TBS-T), incubated for 1 h with secondary IRDye-conjugated antibodies (LI-COR Biosciences) and washed three times in TBS-T for 5 min before near-infrared western blot detection on an Odyssey Imaging System (LI-COR Biosciences).

### Co-immunoprecipitation

A total of 3 × 10^6^ HEK293T cells were plated into 10-cm dishes, cultured for one day, transfected with 9 μg HA-tagged and 9 μg Flag-tagged constructs using TransIT-LT1 transfection reagents (Mirus). The transfected cells were cultured for another two days, treated with 1 μM MLN4924, and co-treated with either degrader or DMSO for 4 h before collection. The cells were collected and lysed in Pierce IP Lysis Buffer (Thermo Fisher Scientific) with Halt Protease Inhibitor Cocktail (Thermo Fisher Scientific) for 20 min on ice, and centrifuged for 15 min to remove the insoluble fraction. Degrader was infused to all buffers used for the degrader-treated arm. For immunoprecipitation, 25 μL of pre-cleaned anti-Flag magnetic beads (Sigma-Aldrich) were added to the lysates. The beads–lysate mix was incubated at 4 °C for 2 h on a rotator. Beads were magnetically removed and washed five times with Pierce IP Lysis Buffer before boiling in 1x NuPAGE LDS Sample Buffer (Thermo Fisher Scientific). Immunoblotting was done as described above.

### Reporter cell line generation

Reporter constructs were generated by BsmBI (New England Biolabs) digestion of Cilantro reporter vector and the insert containing protein of interest coding sequence, followed by ligation with T4 DNA Ligase (New England Biolabs). Constructs were transformed into Stbl3 E. coli and purified using the MiniPrep Kit (Qiagen), and sequences were confirmed by Sanger sequencing (Quintara Biosciences Service). Lentiviruses for reporters were packaged into lentivirus as follows. 0.5 × 10^6^ HEK293T cells were seeded in 2 mL of DMEM media. The next day, a packaging mix including 1.5 μg of psPAX2, 0.15 μg of pVSV-G, and 1.5 μg of transgene plasmid was prepared in 37.5 µL of OptiMEM (Thermo Fisher Scientific). This mix was combined with 9 μL of TransIT-LT1 (Mirus) and 15 µL of OptiMEM, incubated for 30 min at room temperature, and then applied dropwise to cells. Cells were allowed to incubate for another 48 h. Lentivirus was collected by 0.4 μM filters, and then transduced to 2 × 10^6^ of K562-Cas9 or 293T-Cas9 cells at 50% volume ratio by spin infection. One day after infection, reporter cells were selected with puromycin at a concentration of 2 μg/mL.

### Pooled and single-clone knockout cell line generation

sgRNAs targeting DCAF16 (sgDCAF16) or control (sgNTC) were cloned into the sgBFP or sgRFP vector using BsmBI cloning. In brief, vectors were linearized with BsmBI (New England Biolabs) and gel-purified with QIAquick Gel Extraction Kit (Qiagen). Annealed oligos containing sgRNA sequences were phosphorylated with T4 polynucleotide kinase (New England Biolabs) and ligated into linearized vector backbone. sgRNA constructs were transformed, purified, and verified, and lentivirus was generated as described above. Lentivirus containing sgRNA was transduced to 2 × 10^6^ of K562-Cas9 cells at 10% volume ratio by spin infection.

FACS sorting was performed to enrich BFP+ or RFP+ cells one week after infection. For the generation of single-clone DCAF16 knockout cells, pooled K562-Cas9 cells stably expressing sgRNA targeting DCAF16 were seeded in 384-well plates at the density of 0.25 cells per well. Clonal sgDCAF16-expressing K562-Cas9 cells were isolated after one month of expansion, and the genomic sequences were validated via deep sequencing of PCR amplicons targeting sgDCAF16 cutting sites (MGH CCIB DNA Core Service).

### Reporter degradation assays

K562 cells stably expressing degradation reporter were dosed with DMSO or degraders at various time and concentration using D300e Digital Dispenser (HP). The fluorescent signal was quantified by flow cytometry (LSRFortessa flow cytometer, BD Biosciences) and analyzed using FlowJo (flow cytometry analysis software, BD Biosciences). The geometric mean of the eGFP and mCherry fluorescent signal for round and mCherry-positive cells was calculated. GFP expression was normalized to mCherry signal and drug treatments were compared to DMSO controls. The dose-dependent degradation curve was generated using locally weighted smoothing (LOESS) regression in R. The half-maximum degradation concentration (DC50) values of MMH1 and MMH2 were derived using standard four-parameter log-logistic curves fitted with the ‘dr4pl’ R package.

### Competition growth assays

K562-Cas9 cells stably expressing relevant sgRNA with BFP or RFP were mixed with wild-type control cells at 1:9 ratio and plated at 2 × 10^5^ cells per well in a 96 well plate. Cells were dosed with DMSO, 0.1 μM JQ1, 0.33 μM TMX1, or 0.33 μM GNE11 every three to four days. On the same day of drug treatment, cells were split at a 1:3 ratio for maintenance and analyzed by flow cytometry to determine the percentage of BFP+ or RFP+ cells.

### UPS-targeted BRD4_BD2_ reporter CRISPR screen

The Ubiquitin Proteasome System (UPS)-targeted CRISPR library (BISON sgRNA library, addgene 169942^14^) targeting 713 E1, E2 and E3 ubiquitin ligases, deubiquitinases and control genes with a total of 2852 sgRNAs was cloned into the pXPR003 vector. Viruses were produced in a T-175 format as previously described^14^. 2 × 10^6^ of K562-Cas9 or 293T-Cas9 BRD4_BD2_ reporter cell lines were spin infected with BISON virus at 10% volume ratio. Transduced cells were allowed to recover and expand for nine days, then treated with DMSO or degraders. Top (stable gate) and bottom (unstable gate) 5% of cells by eGFP/mCherry fluorescence ratios were sorted for three replicates with at least 1 × 10^5^ cells per replicate. Sorted cells were pelleted, lysed, and sgRNAs were amplified, quantified by next-generation sequencing, and analyzed for enrichment in stable gate over unstable gate, representing degradation rescue.

### Genome-scale and UPS-targeted resistance CRISPR screen

The resistance screen was performed similarly to the BRD4_BD2_ reporter screen with the following modifications. For genome-scale screens, 40 × 10^6^ of K562-Cas9 cells were transduced with viruses generated from genome-wide CRISPR KO Brunello library (addgene 73179^41^) at 10% volume ratio. One day after infection, cells were selected with puromycin at a concentration of 2 μg/mL. Seven days after infection, cells were treated with different compounds or DMSO. The cells were then cultured for 14 more days until collection, with one split every 3–4 days, at which point fresh drug was added. Cells were collected in three replicates, with 2 × 10^6^ cells per replicate, and sgRNAs were isolated and quantified as described above. Results were analyzed by comparing enrichment in the drug-treated arm over the DMSO arm, representing toxicity rescue.

### Data analysis of CRISPR screen

The CRISPR screen data analysis was performed as previously described^14^ and includes the following steps: (1) Reads per sgRNA were normalized to the total number of reads of each sample. (2) For each sgRNA, the enrichment ratio of reads in the stable versus the unstable sorted gate was calculated (for resistance screen, use drug-treated versus DMSO-treated arm), which was then used to rank sgRNAs. (3) The median enrichment ratio of each sgRNA across all sorting or treatment replicates (sgRNA media ratio) was calculated, and the fold change for each gene was determined as the median of sgRNA median ratio of the four sgRNAs targeting the gene. (4) The ranks for each sgRNA were summed for all its replicates, and the gene rank was determined as the median rank of the four sgRNAs targeting the gene. (5) The P values were calculated by simulating a distribution with sgRNAs that had randomly assigned ranks over 100 iterations (two-sided empirical rank-sum test statistics).

### Construction of the BRD4_BD2_ and DCAF16 alanine-scanning library

The BRD4_BD2_ and DCAF16 alanine scanning library constructs were synthesized by Genscript Inc. For the BRD4_BD2_ library, each amino acid of BRD4 between positions 349 and 461 was individually mutated to alanine and each alanine was mutated to arginine. The mutant library was divided into two sub-libraries (_BD2__AlaScan_1/2) and introduced into the Cilantro reporter vector. For the DCAF16 library, each amino acid of DCAF16 from positions 1 and 216 was individually mutated to alanine and each alanine was mutated to arginine. The mutant library was divided into four sub-libraries (DCAF16_AlaScan_1/2/3/4) and introduced into the Ivy-Flag vector.

### BRD4_BD2_ alanine-scanning reporter screen

A total of 2 × 10^6^ K562-Cas9 cells were transduced with BD2_AlaScan_1 or BD2_AlaScan_2 libraries and were selected with 2 μg/mL of puromycin one day after transduction. One week later, cells stably expressing BD2 alanine variant library were treated with DMSO or different BRD4 degraders for 16 h. After treatment, cells were sorted for stable and unstable eGFP/mCherry population, pelleted and lysed using the same method as reporter CRISPR screen. Alanine variant sequences were amplified, quantified by next-generation sequencing, and analyzed for enrichment in the stable gate relative to unstable gate, representing degradation rescue.

### DCAF16 alanine-scanning reporter screen

K562-Cas9 cells with complete DCAF16 knockout were prepared as described above. The DCAF16-KO K562 cells were then transduced with BRD4_BD2_ Cilantro reporter constructs that do not have puromycin selection marker. After reporter construct transduction, mCherry positive cells were sorted to enrich K562 cells stably expressing BRD4_BD2_ reporters. Next, a total of 2 × 10^6^ DCAF16-KO K562 reporter cells were transduced with DCAF16_AlaScan_1, DCAF16_AlaScan_2, DCAF16_AlaScan_3, or DCAF16_AlaScan_4 libraries and selected with puromycin. One week later, cells stably expressing DCAF16 alanine variant library were treated with DMSO or different BRD4 degraders for 16 h. After treatment, cells were sorted, sequencing samples were prepared, and data were analyzed using the same method described above.

### Data analysis of alanine-scanning reporter screen

The alanine scan data analysis was performed as previously described^42^. The analysis pipeline was similar to the above CRISPR screen with the following modifications. (1) The reads of alanine variants, instead of sgRNAs, were used to calculate enrichment ratios and ranks. (2) The read data of each sub-library of the same sorting replicates was concatenated before calculating the ratios and ranks. (3) Only one codon was used for each alanine variants, so the fold change for each alanine variant was determined as the median enrichment ratio of alanine variant across all sorting replicates, and the alanine variant rank was calculated by summing up the ranks across replicates for each alanine variant. Similarly, two-sided empirical rank-sum test was applied to calculate P values.

### Whole-cell proteome mass spectrometry: sample preparation

K562-Cas9 cells were treated at 0.5 μM TMX1 in duplicate, 0.5 μM JQ1 in singlicate, or DMSO control in triplicate for 5 h and harvested by centrifugation before subjected to TMT quantification. In a separate run, K562-Cas9 cells were treated at 0.1 μM MMH1, 0.1 μM MMH2 in duplicates or DMSO control in quadruplicates for 5 h and harvested by centrifugation before subjected to label-free quantification. Cells were lysed by addition of lysis buffer (8 M Urea, 50 mM NaCl, 50 mM 4-(2-hydroxyethyl)-1-piperazineethanesulfonic acid (EPPS) pH 8.5, Protease and Phosphatase inhibitors) and homogenization by bead beating (BioSpec) for three repeats of 30 seconds at 2400. Bradford assay was used to determine the final protein concentration in the clarified cell lysate. 50 µg of protein for each sample was reduced, alkylated and precipitated using methanol/chloroform as previously described^43^, and the resulting washed precipitated protein was allowed to air dry. Precipitated protein was resuspended in 4 M Urea, 50 mM HEPES pH 7.4, followed by dilution to 1 M urea with the addition of 200 mM EPPS, pH 8. Proteins were first digested with LysC (1:50; enzyme:protein) for 12 h at RT. The LysC digestion was diluted to 0.5 M Urea with 200 mM EPPS pH 8 followed by digestion with trypsin (1:50; enzyme:protein) for 6 h at 37 °C.

### Whole-cell proteome mass spectrometry with TMT quantification

Anhydrous ACN was added to each tryptic peptide sample to a final concentration of 30%, followed by addition of Tandem mass tag (TMT) reagents at a labelling ratio of 1:4 peptide:TMT label. TMT labelling occurred over a 1.5 h incubation at room temperature followed by quenching with the addition of hydroxylamine to a final concentration of 0.3%. Each of the samples were combined using adjusted volumes and dried down in a speed vacuum followed by desalting with C18 SPE (Sep-Pak, Waters). The sample was offline fractionated into 96 fractions by high pH reverse-phase HPLC (Agilent LC1260) through an aeris peptide xb-c18 column (phenomenex) with mobile phase A containing 5% acetonitrile and 10 mM NH4HCO3 in LC-MS grade H2O, and mobile phase B containing 90% acetonitrile and 5 mM NH4HCO3 in LC-MS grade H2O (both pH 8.0). The resulting 96 fractions were recombined in a non-contiguous manner into 24 fractions and desalted using C18 solid phase extraction plates (SOLA, Thermo Fisher Scientific) followed by subsequent mass spectrometry analysis. Data were collected using an Orbitrap Fusion Lumos mass spectrometer (Thermo Fisher Scientific, San Jose, CA, USA) coupled with a Proxeon EASY-nLC 1200 LC lump (Thermo Fisher Scientific, San Jose, CA, USA). Peptides were separated on a 50 cm 75 μm inner diameter EasySpray ES903 microcapillary column (Thermo Fisher Scientific). Peptides were separated over a 190 min gradient of 6 - 27% acetonitrile in 1.0% formic acid with a flow rate of 300 nL/min.

Quantification was performed using a MS3-based TMT method as described previously^44^. The data were acquired using a mass range of m/z 340 – 1350, resolution 120,000, AGC target 5 × 10^5^, maximum injection time 100 ms, dynamic exclusion of 120 seconds for the peptide measurements in the Orbitrap. Data dependent MS2 spectra were acquired in the ion trap with a normalized collision energy (NCE) set at 35%, AGC target set to 1.8 × 10^4^ and a maximum injection time of 120 ms. MS3 scans were acquired in the Orbitrap with HCD collision energy set to 55%, AGC target set to 2 × 10^5^, maximum injection time of 150 ms, resolution at 50,000 and with a maximum synchronous precursor selection (SPS) precursors set to 10.

### LC-MS data analysis for TMT quantification

Proteome Discoverer 2.4 (Thermo Fisher Scientific) was used for .RAW file processing and controlling peptide and protein level false discovery rates, assembling proteins from peptides, and protein quantification from peptides. The MS/MS spectra were searched against a Swissprot human database (January 2021) containing both the forward and reverse sequences. Searches were performed using a 10 ppm precursor mass tolerance, 0.6 Da fragment ion mass tolerance, tryptic peptides containing a maximum of two missed cleavages, static alkylation of cysteine (57.0215 Da), static TMT labelling of lysine residues and N-termini of peptides (229.1629), and variable oxidation of methionine (15.9949 Da). TMT reporter ion intensities were measured using a 0.003 Da window around the theoretical m/z for each reporter ion in the MS3 scan. The peptide spectral matches with poor quality MS3 spectra were excluded from quantitation (summed signal-to-noise across channels < 100 and precursor isolation specificity < 0.5), and the resulting data was filtered to only include proteins with a minimum of 2 unique peptides quantified. Reporter ion intensities were normalized and scaled using in-house scripts in the R framework (R Core Team, 2014). Significant changes comparing the relative protein abundance between samples were assessed by two-side moderated t-test as implemented in the limma package within the R framework^45^.

### Whole-cell proteome mass spectrometry with label free quantification

Sample digests were acidified with formic acid to a pH of 2-3 prior to desalting using C18 solid phase extraction plates (SOLA, Thermo Fisher Scientific). Desalted peptides were dried in a vacuum-centrifuged and reconstituted in 0.1% formic acid for LC-MS analysis.

Data were collected using a TimsTOF Pro2 (Bruker Daltonics, Bremen, Germany) coupled to a nanoElute LC pump (Bruker Daltonics, Bremen, Germany) via a CaptiveSpray nano-electrospray source. Peptides were separated on a reversed-phase C18 column (25 cm × 75 μM ID, 1.6 μM, IonOpticks, Australia) containing an integrated captive spray emitter. Peptides were separated using a 50 min gradient of 2 - 30% buffer B (acetonitrile in 0.1% formic acid) with a flow rate of 250 nL/min and column temperature maintained at 50 ºC.

DDA was performed in Parallel Accumulation-Serial Fragmentation (PASEF) mode to determine effective ion mobility windows for downstream diaPASEF data collection^46^. The ddaPASEF parameters included: 100% duty cycle using accumulation and ramp times of 50 ms each, 1 TIMS-MS scan and 10 PASEF ramps per acquisition cycle. The TIMS-MS survey scan was acquired between 100 – 1700 m/z and 1/k0 of 0.7 - 1.3 V.s/cm2. Precursors with 1 – 5 charges were selected and those that reached an intensity threshold of 20,000 arbitrary units were actively excluded for 0.4 min. The quadrupole isolation width was set to 2 m/z for m/z <700 and 3 m/z for m/z >800, with the m/z between 700-800 m/z being interpolated linearly. The TIMS elution voltages were calibrated linearly with three points (Agilent ESI-L Tuning Mix Ions; 622, 922, 1,222 m/z) to determine the reduced ion mobility coefficients (1/K0). To perform diaPASEF, the precursor distribution in the DDA m/z-ion mobility plane was used to design an acquisition scheme for DIA data collection which included two windows in each 50 ms diaPASEF scan. Data was acquired using sixteen of these 25 Da precursor double window scans (creating 32 windows) which covered the diagonal scan line for doubly and triply charged precursors, with singly charged precursors able to be excluded by their position in the m/z-ion mobility plane. These precursor isolation windows were defined between 400 - 1200 m/z and 1/k0 of 0.7 - 1.3 V.s/cm2.

### LC-MS data analysis for label free quantification

The diaPASEF raw file processing and controlling peptide and protein level false discovery rates, assembling proteins from peptides, and protein quantification from peptides was performed using library free analysis in DIA-NN 1.8^47^. Library free mode performs an in-silico digestion of a given protein sequence database alongside deep learning-based predictions to extract the DIA precursor data into a collection of MS2 spectra. The search results are then used to generate a spectral library which is then employed for the targeted analysis of the DIA data searched against a Swissprot human database (January 2021). Database search criteria largely followed the default settings for directDIA including tryptic with two missed cleavages, carbomidomethylation of cysteine, and oxidation of methionine and precursor Q-value (FDR) cut-off of 0.01. Precursor quantification strategy was set to Robust LC (high accuracy) with RT-dependent cross run normalization. Proteins with missing values in any of the treatments and with poor quality data were excluded from further analysis (summed abundance across channels of <100 and mean number of precursors used for quantification <2). Protein abundances were scaled using in-house scripts in the R framework (R Development Core Team, 2014). Significant changes comparing the relative protein abundance between samples were assessed by two-side moderated t-test as implemented in the limma package within the R framework^45^.

### Immunoprecipitation mass spectrometry (IP-MS)

For IP-MS experiments, immunoprecipitation (IP) was performed as described above. After the washing step, samples were eluted using Glycine-HCl buffer (0.2 M, pH 2.4). The IP eluates were reduced with 10 mM TCEP for 30 min at room temperature, and then alkylated with 15 mM iodoacetamide for 45 min at room temperature in the dark. Alkylation was quenched by the addition of 10 mM DTT. Proteins were isolated by methanol-chloroform precipitation. The protein pellets were dried and then resuspended in 50 μL 200 mM EPPS pH 8.0. The resuspended protein samples were digested with 2 μg LysC overnight at room temperature followed by the addition of 0.5 μg Trypsin for 6 h at 37°C. Protein digests were dried, resuspended in 100 μL 1% formic acid, and desalted using 10-layer C18 stage-tips before being analyzed by LC-MS.

Data were collected using an Orbitrap Exploris 480 mass spectrometer (Thermo Fisher Scientific) equipped with a FAIMS Pro Interface and coupled with a UltiMate 3000 RSLCnano System. Peptides were separated on an Aurora 25 cm × 75 μm inner diameter microcapillary column (IonOpticks), and using a 60 min gradient of 5 - 25% acetonitrile in 1.0% formic acid with a flow rate of 250 nL/min.

Each analysis used a TopN data-dependent method. The FAIMS Pro Interface compensation voltages were set to -50 and -70. The data were acquired using a mass range of m/z 350 – 1200, resolution 60,000, AGC target 3 × 10^6^, auto maximum injection time, dynamic exclusion of 15 sec, and charge states of 2-6. TopN 20 data-dependent MS2 spectra were acquired with a scan range starting at m/z 110, resolution 15,000, isolation window of 1.4 m/z, normalized collision energy (NCE) set at 30%, AGC target 1 × 10^5^ and the automatic maximum injection time.

### LC-MS data analysis for IP-MS

Proteome Discoverer 2.4 (Thermo Fisher Scientific) was used for .RAW file processing and controlling peptide and protein level false discovery rates, assembling proteins from peptides, and protein quantification from peptides. MS/MS spectra were searched against a Uniprot human database (January 2021) with both the forward and reverse sequences as well as known contaminants such as human keratins. Database search criteria were as follows: tryptic with two missed cleavages, a precursor mass tolerance of 10 ppm, fragment ion mass tolerance of 0.6 Da, static alkylation of cysteine (57.02146 Da) and variable oxidation of methionine (15.99491 Da). Peptides were quantified using the MS1 Intensity, and peptide abundance values were summed to yield the protein abundance values.

Resulting data was filtered to only include proteins that had a minimum of 2 abundance counts in at least two runs. Abundances were normalized and scaled using in-house scripts in the R framework. Missing values in the dataset were imputed by random selection from a gaussian distribution centered around the mean of the existing data and with the mean relative standard deviation of the dataset. Significant changes comparing the relative protein abundance between samples were assessed by two-side moderated t-test as implemented in the limma package within the R framework^45^. A protein was considered a ‘hit’ if it met our predetermined ‘hit’ threshold of P-value < 0.01 and fold change > 2.

### Protein expression and purification

The human wild-type BRD4_BD1_ and BRD4_BD2_ (UniProt entry O60885, residues 75-147 and 368-440) were subcloned into E.Coli pET100/D-TOPO vector with N-terminal His6-Avi fusions, and expressed in E. Coli BL21-DE3 Rosetta cells using standard protocols. Biotinylation of BRD4_BD1_ and BRD4_BD2_ was done as previously described^44^.

The human wild-type and mutant versions of DCAF16 (UniProt entry Q9NXF7, full length), DDB1ΔB (Uniport entry Q16531, residues 396–705 replaced with GNGNSG linker), and DDA1 (Uniport entry Q9BW61, full length) were cloned in pAC-derived vectors^48^, and recombinant proteins were expressed as N-terminal His6 (DDA1), StrepII-Avi (DCAF16) or His6-3C-Spy (DDB1ΔB) fusions in Trichoplusia ni High-Five insect cells using the baculovirus expression system (Invitrogen). Briefly, expression plasmids were transfected into Spodoptera frugiperda (Sf9) cells at a density of 0.9 × 10^6^ cells/mL grown in ESF 921 medium (Expression Systems) to generate baculovirus, and this was followed by two rounds of infection in Sf9 cells to increase viral titer. For recombinant protein expression, Hi Five cells grown in Sf-900 II SFM media (Gibco) at a density of 2.0 × 10^6^ cells/mL were infected with baculovirus at 1.5% v/v ratio. After 40 h of expression at 27 ºC, Hi Five cells were collected by centrifugation for 15 min at 3,500 × g.

For purification of StrepII or His6-tagged proteins, pelleted cells were resuspended in lysis buffer containing 50 mM tris (hydroxymethyl) aminomethane hydrochloride (Tris–HCl) pH 8.0, 200 mM NaCl, 2 mM tris (2-carboxyethyl) phosphine (TCEP), 1 mM phenylmethylsulfonyl fluoride (PMSF), and protease inhibitors, and the cell pellets were lysed by sonication. Following ultracentrifugation (1h, 185,511 x g), the soluble fraction was passed over the appropriate affinity resin of Strep-Tactin XT Superflow (IBA) or Ni Sepharose 6 Fast Flow affinity resin (GE Healthcare), eluted with wash buffer (50 mM Tris–HCl pH 8.0, 200 mM NaCl, 1 mM TCEP) supplemented with 50 mM d-Biotin (IBA) or 100 mM imidazole (Fisher Chemical), respectively. The affinity-purified proteins were then applied to an ion exchange column (Poros 50HQ) and eluted in 50 mM Tris-HCl pH 8.5 and 2 mM TCEP by a linear salt gradient (from 50–800 mM NaCl). Purified DCAF16 was dephosphorylated with lambda-phosphatase (NEB) at 4 ºC overnight. Both DCAF16 complex and BRD4_BD2_ were cleaved with TEV protease at 4 ºC overnight. They were then subjected to size-exclusion chromatography (SEC) on a Superdex 200 Increase 10/300 (GE Healthcare) in 50 mM 4-(2-hydroxyethyl)-1-piperazineethanesulfonic acid (HEPES) pH 7.4 or pH 8.0, 200 mM NaCl and 2 mM TCEP. Peak fractions were pooled, concentrated, flash-frozen in liquid nitrogen, and stored at −80 ºC.

### BODIPY-FL-Spycatcher labeling of DCAF16-DDB1ΔB

Purified StrepII-Avi-DCAF16 + His6-3C-Spy-DDB1ΔB was incubated overnight at 4°C with BODIPY-FL labeled SpyCatcherS50C protein at stoichiometric ratio. Protein was concentrated and loaded on the Enrich SEC 650 10/300 (Bio-rad) size exclusion column and the labeling was monitored with absorption at 280 and 490 nm. The protein peak corresponding to the labeled protein was pooled, concentrated by ultrafiltration (Millipore), and flash frozen.

### DDB1-DCAF16–BRD4BD TR-FRET

Titrations of compounds to induce the DCAF16-BRD4BD complex were carried out by mixing 100 nM biotinylated BRD4_BD1_ or BRD4_BD2_, 500 nM BODIPY-FL labeled DDB1ΔB-DCAF16 variants, and 2 nM terbium-coupled streptavidin (Invitrogen) in an assay buffer containing 50 mM HEPES pH 8.0, 200 mM NaCl, 0.1% Pluronic F-68 solution (Sigma), 0.5% bovine serum albumin (BSA) (w/v) and 1 mM TCEP. After dispensing the assay mixture (15 μL volume), increasing concentrations of compounds were dispensed in a 384-well microplate (Corning, 4514) using a D300e Digital Dispenser (HP) normalized to 1% DMSO. After excitation of terbium fluorescence at 337 nm, emission at 490 nm (terbium) and 520 nm (BODIPY FL) were recorded with a 70 μs delay over 600 μs to reduce background fluorescence, and the reaction was followed over 60 cycles of each data point using a PHERAstar FS microplate reader (BMG Labtech). The TR-FRET signal of each data point was extracted by calculating the 520/490 nm ratio. The dose-dependent TR-FRET curve was generated using locally weighted smoothing (LOESS) regression in R.

### Intact protein mass spectrometry

Prior to intact mass analysis, recombinant human DDB1ΔB-DCAF16 variants were incubated with DMSO, TMX1, KB02-JQ1, MMH1, or MMH2 with and without the presence of recombinant human BRD4_BD2_ for 16 h at 4°C. For GNE11, recombinant proteins were incubated with drug at room temperature for 16 h. Intact mass analysis of DCAF16 variants was performed similarly to a previously described protocol^49^ with modifications. Briefly, drug-treated proteins were injected on a self-packed column (6 cm POROS 50R2 packed in 0.5 mm I.D. tubing), desalted for 4 minutes, and then eluted to an LTQ ion trap mass spectrometer (Thermo Fisher Scientific) using an HPLC gradient (0-100% B in 20 minutes, A=0.1M acetic acid, B=0.1 M acetic acid in acetonitrile, ESI spray voltage=5kV). The mass spectrometer acquired full scan mass spectra (m/z 300-2000) in profile mode. Mass spectra were deconvoluted using MagTran version 1.03 b2^50^. Labeling efficiency was calculated from zero charge mass spectra using peak heights according to [peak height labeled protein] / [peak height labeled protein + peak height unlabeled protein] x 100%.

### EM Sample Preparation and data collection

The ternary complex was incubated at RT for 30 min at 15 μM DCAF16-DDB1ΔB-DDA1 complex, 25 μM BRD4_BD2_, and 50 μM MMH2 before loading on a Superdex 200 Increase 10/300 SEC column. After SEC, the purified DCAF16 complex was incubated with an extra 1.2x molar excess of purified BRD4_BD2_ for 30 minutes at 4 ºC and then mixed with 0.011% Lauryl Maltose Neopentyl Glycol (LMNG) right before preparation of cryo-EM grids. Glow-discharged Quantifoil UltrAuFoil 0.6/1.0 grids were prepared using a Leica EM-GP, operated at 10°C and 90% relative humidity. 4 μL of sample (1.25 mg/mL) were applied, incubated on the grid for 10 s, and blotted for 3 sec before vitrification. Grids were imaged in a Titan Krios equipped with a Gatan Quantum Image filter (20 eV slit width) and a post-GIF Gatan K3 direct electron detector. 17,118 movies were acquired at 300 kV at a nominal magnification of 105,000 x in counting mode with a pixel size of 0.83 Å/pixel using SerialEM ver. 4.0.5^51^. One movie (40 frames each) was acquired per hole with nine holes per stage position (resulting in 9 image acquisition groups), in a defocus range from −0.8 - −2.0 μm over an exposure time of 2.30 s and a total dose of 50.27 e/Å^2^.

### EM Data Processing and model building

All processing was performed in cryoSPARCv3.3.2^52^. 17,118 movies were corrected for beam-induced motion and contrast transfer function was estimated on-the-fly in cryoSPARC live. 14,452,363 particles were extracted (at 1.58 Å/pixel) from 15,448 curated micrographs after TOPAZ particle picking^53^. The extracted particles were split into two batches and sent through two rounds of 2D classification to remove mispicks and DDB1-only classes. The resulting particles were combined and further cleaned by an additional round of 2D classification. The remaining 4,795,088 particles were classified by 3D variability^54^ in clustering mode (8 clusters). Particles from 3 clusters (1,433,050) with most pronounced density for BRD4_BD2_ were combined, re-extracted at 0.89 Å/pixel and a final homogeneous refinement was followed by local refinement using a mask encompassing the whole particle. The final reconstruction reached a resolution of 2.2 Å, based on the Fourier shell correlation (FSC) 0.143 threshold criterion^55,56^. This map, sharpened with a *B*-value of -74.4 Å^2^, as well as a map post-processed using deepEMhancer^40^ were used for model building in COOTv0.9.8^57^. Models for DDB1, DDA1 (pdb: 6Q0R^58^) and BRD4_BD2_ (pdb: 6VIX^59^) were first fit as rigid bodies in ChimeraXv1.4^60^, relaxed into the density using ISOLDEv1.3^61^, and then adjusted manually in COOT. The model for DCAF16 was built de novo. A component dictionary for the MMH2 compound and a link dictionary for MMH2 linked to cysteine were generated using AceDRG^62,63^. The compound was linked to _Cys58_ and the model was refined iteratively in Refmac5^64^ and phenix.real_space_refine^65,66^ (v.1.19.2-4158). The resulting model was deposited in the PDB under accession code 8G46. The final map was deposited as main map in the EMDB (EMD-29714) with the map from deepEMhancer as additional map. Interface areas were calculated using PDBePisa^30^, structural similarity searches were conducted using PDBeFold^67^, and all figures with models and density were generated in ChimeraX (v2.5.4, Schrödinger LLC). The local resolution range is given based on the 0-75% percentile in local resolution histograms^68^. Directional resolution was calculated using 3DFSC^69^. Structural biology applications used in this project were compiled and configured by SBGrid^70^.

### BRD4_BD1_ and BRD4_BD2_ AlphaScreen assays

The AlphaScreen assays were performed with minor modifications from the manufacturer’s protocol (Perkin Elmer). All reagents were diluted in AlphaScreenTM buffer (50 mM HEPES, 150 mM NaCl, 0.01 % v/v Tween-20, 0.1 % w/v BSA, pH 7.4). After addition of the Alpha beads to the master solutions, all subsequent steps were performed under low light conditions. A 2x solution of components with final concentrations of His-BRD4_BD1_, His-BRD4_BD2_ at 20 nM, Ni-coated acceptor bead at 10 μg/mL, and biotinylated-JQ1 at 10 nM was added in 10 μL to 384-well plates (AlphaPlate-384, PerkinElmer). Plates were spun down at 1000 rpm. A 10-point serial dilution of compounds in DMSO was prepared at 200x of the final concentration.

Compound (100 nL) from these stock plates was added by pin transfer using a Janus Workstation (PerkinElmer). A 2x solution of streptavidin-coated donor beads with a final concentration of 10 μg/mL was added in a 10 μL volume. The plates were spun down again at 1000 rpm and sealed with foil to prevent light exposure and evaporation. The plates were then incubated at room temperature for 1 h and read on an Envision 2104 (PerkinElmer) using the manufacturer’s protocol. After normalization to DMSO-treated negative control wells, the dose-dependent activity inhibition curve was generated using standard four-parameter log-logistic curves fitted with the ‘dr4pl’ R package.

## Data Availability

Cryo-EM maps and coordinates have been deposited in the EMDB and PDB, under accession codes EMD-29714 and 8G46 respectively. Synthetic procedures of JQ1-derived compounds are provided in Supplementary Note. Proteomics data sets will be deposited in PRIDE before publication. Uncropped western blots, deep sequencing data for DCAF16 knockout clones, and flow cytometry gating strategy will be provided as Supplementary Figures before publication. Primary data for CRISPR screens and alanine-scanning mutagenesis screens will be provided as Supplementary Data before publication.

## Code Availability

Codes used to generate dose-response curves, volcano plots, and correlation plots will be provided as Supplementary Codes before publication.

## Acknowledgements

We thank the Broad Institute Walk-Up Sequencing team, the Broad Institute Genetic Perturbation Platform, and the Broad Institute PRISM team for technical assistance. We thank the staff of Harvard Cryo-Electron Microscopy Center for Structural Biology for their technical expertise and support during grid screening and data collection. We acknowledge the SBGrid consortium for assistance with structural biology software packages. We thank members of the Eck laboratory for valuable structural discussions. We are grateful to all members of the Ebert, Fischer, and Gray laboratories for discussions on many project-related topics. Y.L. was supported by Harvard Institutional Stipend and the Genevieve Castrodale Carpenter Graduate Financial Aid Fund. M.W.M. was supported by the Chleck Fellowship Foundation and the Fujifilm Fellowship. M.T. was a CPRIT scholar in cancer research was supported by the CPRIT research funding (RR220012). This work was supported by the National Institutes of Health (NIH) grants R01HL082945, P01CA066996, P50CA206963, and R35CA253125 (to B.L.E.), and the Howard Hughes Medical Institute (to B.L.E.), NIH grants R01CA262188 and P01CA066996 (to E.S.F.), and the Mark Foundation for Cancer Research 19-001-ELA (to E.S.F.), NIH High End Instrumentation grant (1S10OD028697-01) (to N.S.G.), and the departmental funds from Stanford Chemical and Systems Biology and Stanford Cancer Institute (to N.S.G.), NIH grants U24DK116204, R01CA219850, R01CA233800, R21CA247671 (to J.A.M.), and the Mark Foundation for Cancer Research, the Massachusetts Life Science Center (to J.A.M.).

## Author Contributions

Y.L., B.L.E., E.S.F. and N.S.G. conceptualized and initiated the study; Y.L. designed and performed functional genomics studies, mutagenesis screens, and cellular validation experiments with the help of B.S., S.X., C.Z., J.M.T., P.M.C.P., H.Y., and M.S.; M.W.M. designed and carried out biochemical studies and structural analyses with the help of M.H., K.P., C.Y.J., and R.P.N.; M.M.H. and M.T. developed and synthesized covalent BRD4 molecular glue degraders with the help of B.J.G. and F.C.M.; R.L. and K.A.D. performed whole-cell proteomics and IP-MS experiments; S.B.F performed intact mass spectrometry experiments with the help of I.T.; M.Y.W. performed AlphaScreen experiments with the help of L.H.S.; B.L.E., E.S.F., N.S.G., J.A.M., and J.Q. supervised the project. Y.L., M.W.M., M.M.H., B.L.E., E.S.F. and N.S.G. wrote the manuscript with input from all authors.

## Additional Information

Supplementary Information is available for this paper.

Correspondence and requests for materials should be addressed to B.L.E., E.S.F., and N.S.G.

## Extended Data Figures and Tables

**Extended Data Figure 1.**
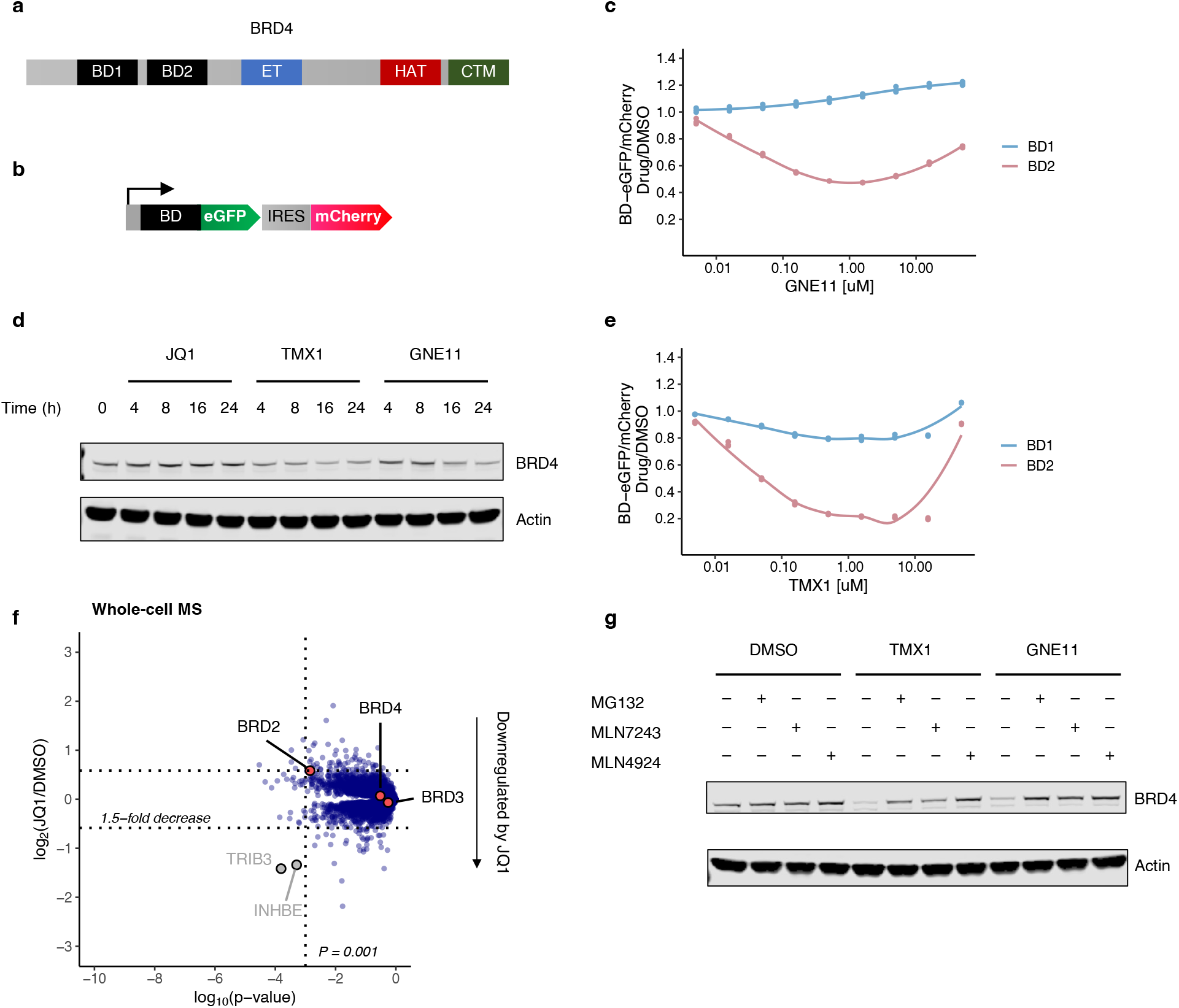
Degradation characterization of JQ1-derived compounds. **a**. The domain structure of BRD4. **b**. Schematic of BRD4BD stability reporter. IRES, internal ribosome entry site. **c**. Flow cytometry analysis of BRD4_BD1_-eGFP and BRD4_BD2_-eGFP degradation in K562 cells that were treated with increasing concentrations of GNE11 for 16 h (n=3). **d**. Western blots of BRD4 degradation in K562 cells that were treated with JQ1, TMX1 or GNE11 at 1 μM for increasing time points. **e**. Flow cytometry analysis of BRD4_BD1_-eGFP and BRD4_BD2_-eGFP degradation in K562 cells that were treated with increasing concentrations of TMX1 for 16 h (n=3). **f**. Quantitative whole proteome analysis of K562 cells after treatment with JQ1 at 0.5 μM (n=1) or DMSO (n=3) for 5 h. **g**. Western blots of BRD4 degradation in K562 cells that were treated with DMSO, TMX1 at 1 μM, GNE11 at 1 μM, MG132 at 10 μM, MLN7243 at 1 μM, and MLN4924 at 1 μM for 16 h.

**Extended Data Figure 2.**
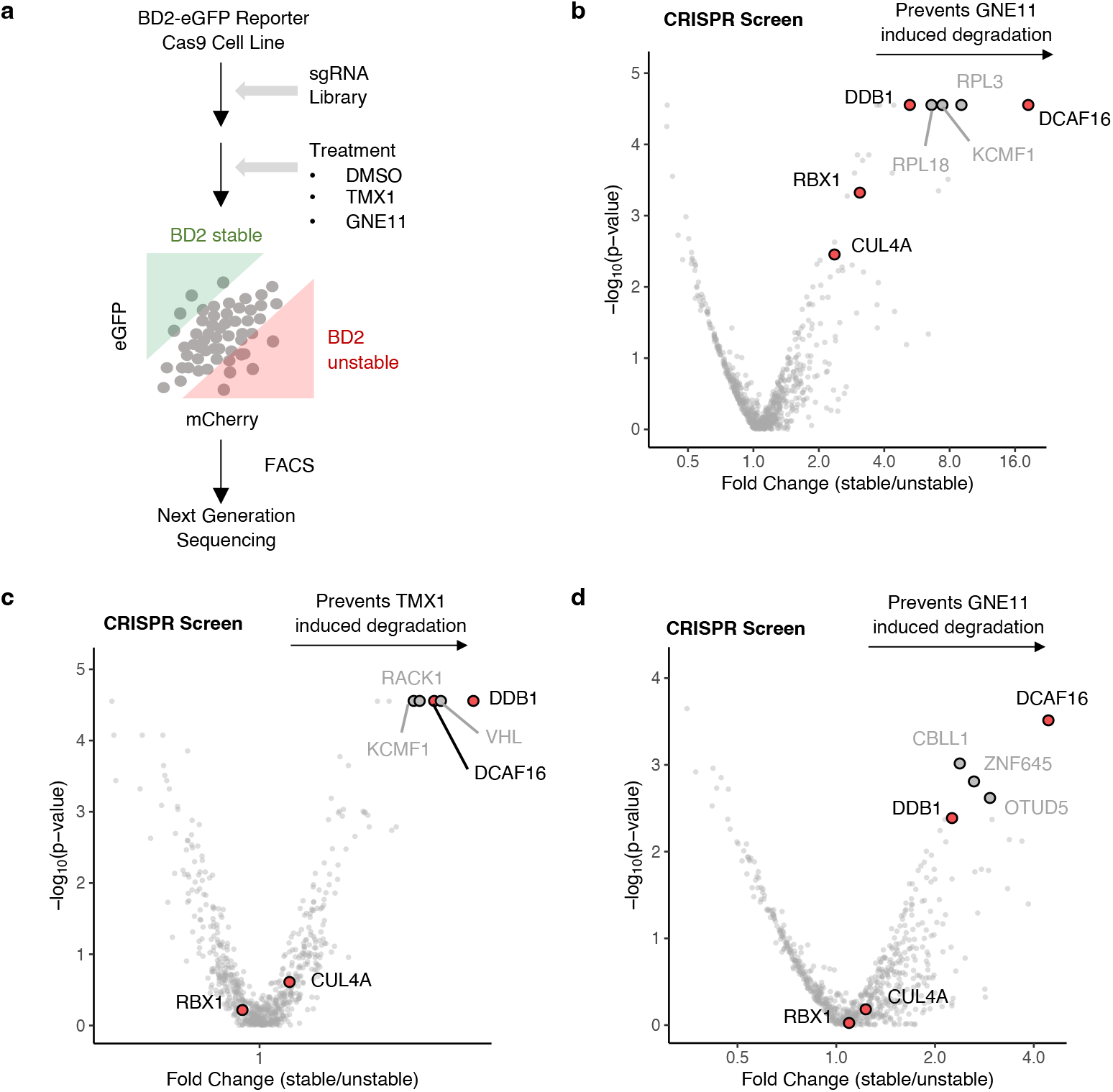
UPS-targeted BRD4_BD2_ reporter CRISPR screen for JQ1-derived compounds. **a**. Schematic of the CRISPR degradation screen for BRD4_BD2_ stability. **b**. UPS-focused CRISPR degradation screen for BRD4_BD2_-eGFP stability in K562-Cas9 cells treated with GNE11 at 1 μM for 16 h (n=2). **c**. UPS-focused CRISPR degradation screen for BRD4_BD2_-eGFP stability in 293T-Cas9 cells treated with TMX1 at 1 μM for 16 h (n=2). **d**. UPS-focused CRISPR degradation screen for BRD4_BD2_-eGFP stability in 293T-Cas9 cells treated with GNE11 at 1 μM for 16 h (n=2).

**Extended Data Figure 3.**
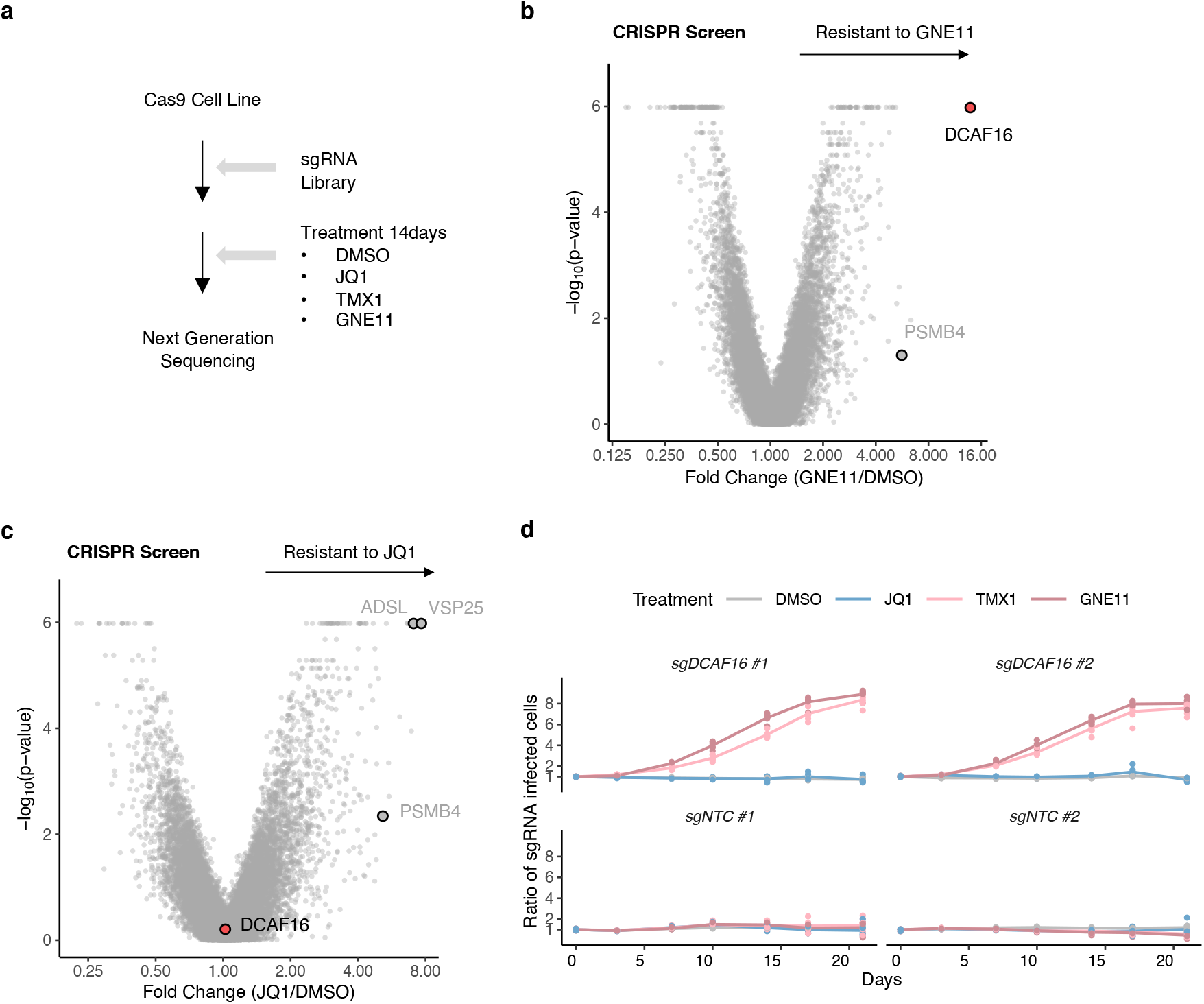
Genome-scale resistance CRISPR screen for JQ1-derived compounds. **a**. Schematic of the CRISPR resistance screen. **b**. Genome-wide CRISPR resistance screen in K562-Cas9 cells treated with GNE11 at 0.1 μM (n=3) or DMSO (n=3) for 14 days. **c**. Genome-wide CRISPR resistance screen in K562-Cas9 cells treated with JQ1 at 0.1 μM (n=3) or DMSO (n=3) for 14 days. **d**. Flow cytometry-based competitive growth assay of K562-Cas9 cells expressing BFP- or RFP-tagged sgRNAs against DCAF16 and non-targeting control (NTC) treated with DMSO, JQ1 at 0.1 μM, TMX1 at 0.1 μM, or GNE11 at 0.1 μM for increasing time points (n=3).

**Extended Data Figure 4.**
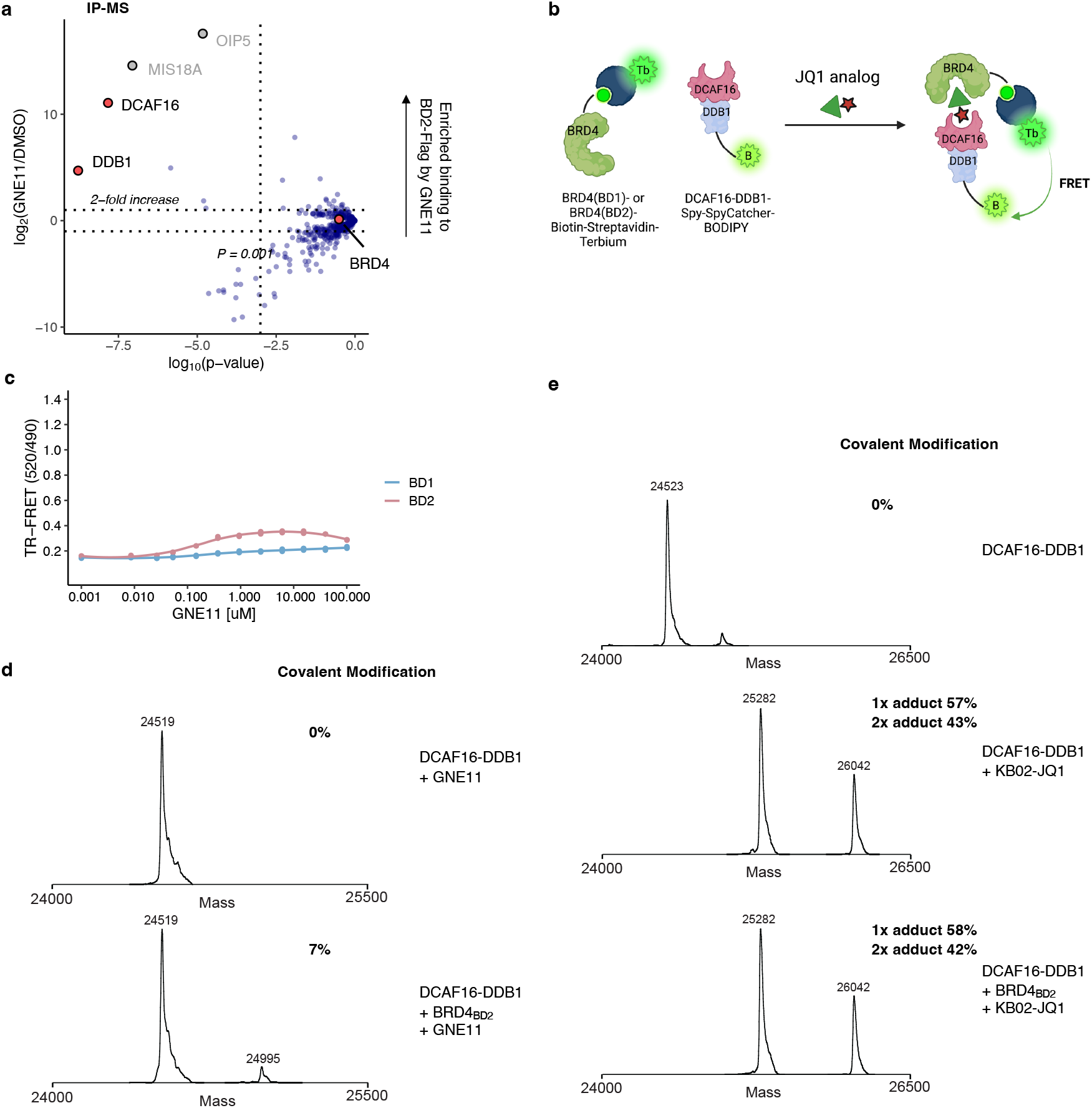
Covalent recruitment of DCAF16 to BRD4_BD2_. **a**. Flag immunoprecipitation (IP) followed by mass spectrometry in 293T cells overexpressing BRD4_BD2_-Flag of cells treated with either MLN4924 plus GNE11 both at 1 μM (n=4), or MLN4924 at 1 μM only (n=4). Fold enrichment and p-values were calculated by comparing GNE11/MLN4924 treated samples to MLN4924 only control samples. **b**. Schematic of the TR-FRET set-up. Positions of FRET donor (terbium-coupled streptavidin) and acceptor (BODIPY– SpyCatcher) are indicated in the structural model. **c**. TR-FRET signal for DDB1-DCAF16-BODIPY to BRD4BD1-terbium or BRD4_BD2_-terbium with increasing concentrations of GNE11 (n=3). **d**. Intact protein mass spectra of DDB1-DCAF16 co-incubated with GNE11 at 25°C for 16 h, or DDB1-DCAF16 co-incubated with GNE11 and BRD4_BD2_ at 25°C for 16 h. **e**. Intact protein mass spectra of DDB1-DCAF16 alone, DDB1-DCAF16 co-incubated with KB02-JQ1 at 4°C for 16 h, or DDB1-DCAF16 co-incubated with KB02-JQ1 and BRD4_BD2_ at 4°C for 16 h.

**Extended Data Figure 5.**
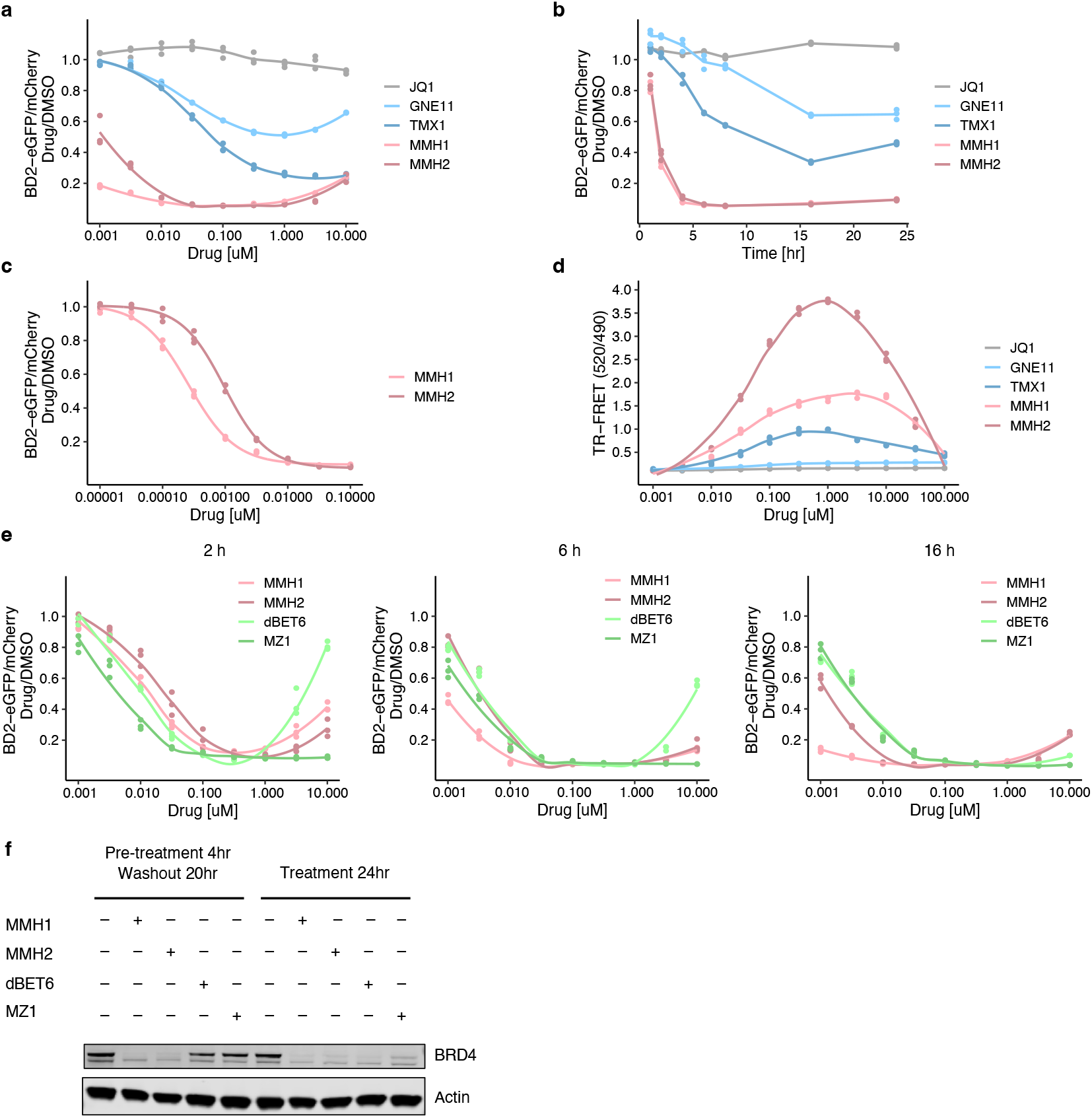
Optimized electrophilic warheads increases potency of degraders. **a**. Flow cytometry analysis of BRD4_BD2_-eGFP degradation in K562 cells that were treated with increasing concentrations of JQ1, GNE11, TMX1, MMH1 or MMH2 for 16 h (n=3). **b**. Flow analysis of BRD4_BD2_-eGFP degradation in K562 cells that were treated with JQ1 at 1 μM, TMX1 at 1 μM, GNE11 at 1 μM, MMH1 at 0.1 μM or MMH2 at 0.1 μM for increasing time points (n=3). **c**. Flow cytometry analysis of BRD4_BD2_-eGFP degradation in K562 cells that were treated with increasing concentrations of MMH1 or MMH2 for 16 h (n=3). **d**. TR-FRET signal for DDB1-DCAF16-BODIPY to BRD4_BD2_-terbium with increasing concentrations of JQ1, GNE11, TMX1, MMH1 or MMH2 (n=3). **e**. Flow cytometry analysis of BRD4_BD2_-eGFP degradation in K562 cells that were treated with increasing concentrations of MMH1, MMH2, dBET6 or MZ1 for 2 h, 6 h, or 16 h (n=3). **f**. Western blot of BRD4 degradation in K562 cells pre-treated with MMH1, MMH2, dBET6 or MZ1 at 0.1 μM for 4 h, washed with PBS and resuspended in fresh media or the same drug-treated media for an additional 20 h.

**Extended Data Figure 6.**
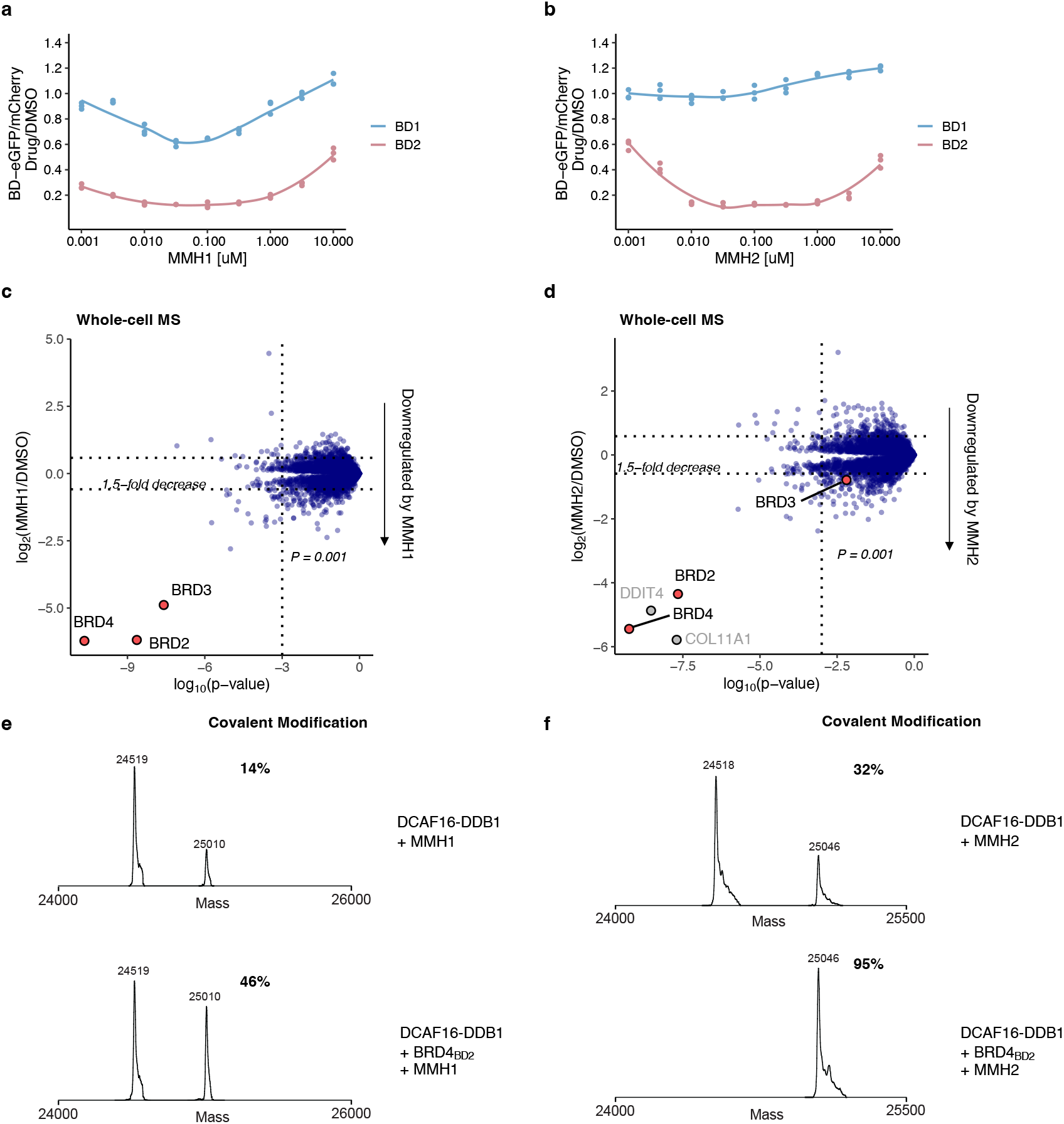
MMH1 and MMH2 conserve the mechanism of action as TMX1 and GNE11. **a**. Flow cytometry analysis of BRD4_BD1_-eGFP and BRD4_BD2_-eGFP degradation in K562 cells that were treated with increasing concentrations of MMH1 for 16 h (n=3). **b**. Flow cytometry analysis of BRD4_BD1_-eGFP and BRD4_BD2_-eGFP degradation in K562 cells that were treated with increasing concentrations of MMH2 for 16 h (n=3). **c**. Quantitative whole proteome analysis of K562 cells after treatment with MMH1 at 0.1 μM (n=2) or DMSO (n=4) for 5 h. **d**. Quantitative whole proteome analysis of K562 cells after treatment with MMH2 at 0.1 μM (n=2) or DMSO (n=4) for 5 h. **e**. Intact protein mass spectra of DDB1-DCAF16 co-incubated with MMH1 at 4°C for 16 h, or DDB1-DCAF16 co-incubated with MMH1 and BRD4_BD2_ at 4°C for 16 h. **f**. Intact protein mass spectra of DDB1-DCAF16 co-incubated with MMH2 at 4°C for 16 h, or DDB1-DCAF16 co-incubated with MMH2 and BRD4_BD2_ at 4°C for 16 h.

**Extended Data Figure 7.**
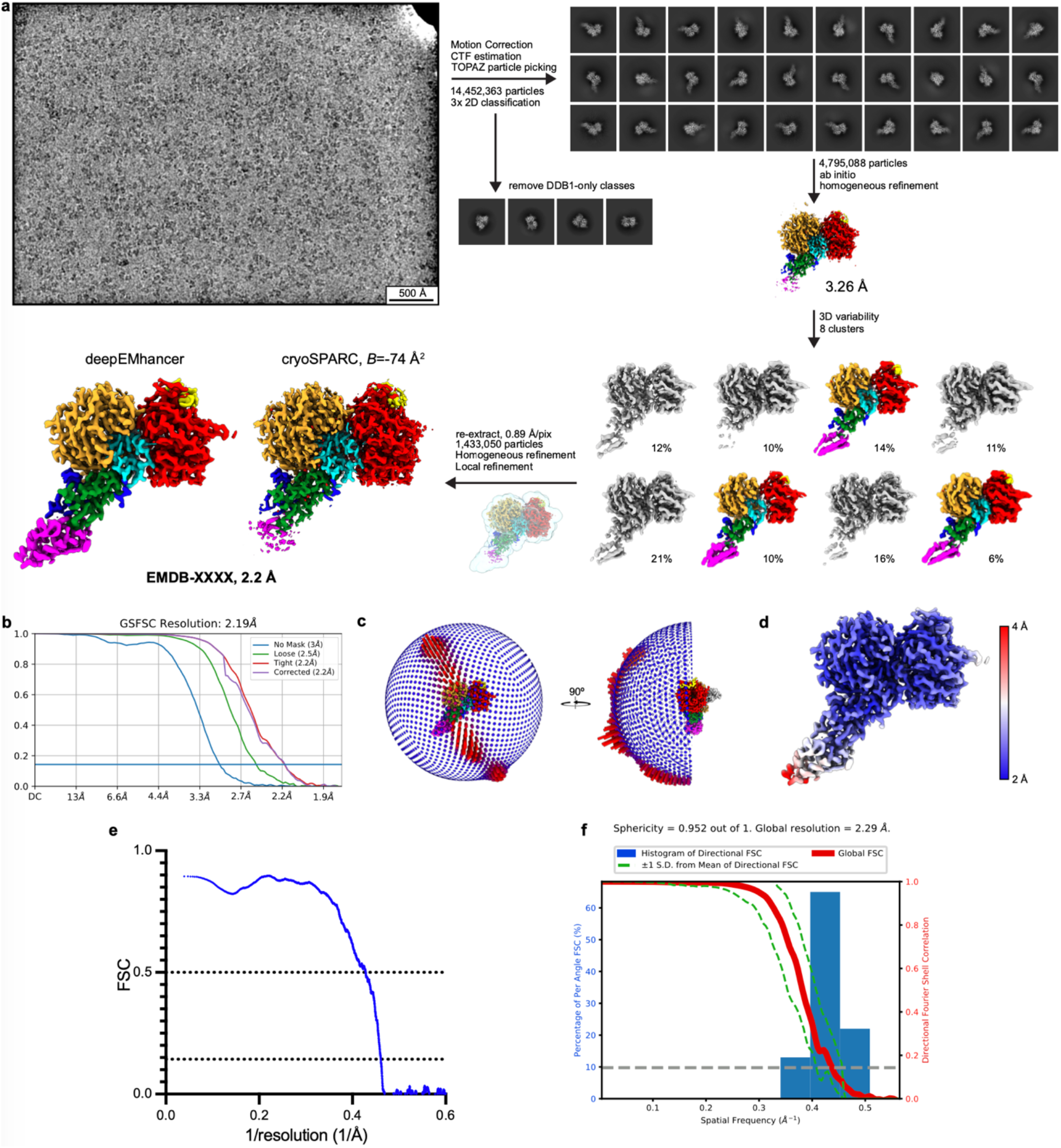
Cryo-EM processing workflow of DDB1ΔB-DDA1-DCAF16 in complex with BRD4_BD2_ and MMH2. **a**. Overview of processing workflow for the DDB1**Δ**B-DDA1-DCAF16-BRD4_BD2_-MMH2 dataset, from raw micrographs (low pass-filtered to 5 Å) to final maps. All steps performed in cryoSPARC^52^. Particles belonging to colored volumes were taken into the subsequent steps. Maps here and in following panels (unless noted otherwise) are contoured at 0.3 (clusters), 0.743 (initial consensus), 0.6 (final from cryoSPARC), 0.2 (final from deepEMhancer). **b**. FSC plot. **c**. Viewing distribution for the final reconstruction. **d**. Local resolution mapped onto final map. **e**. Model-to-map FSC, dotted lines indicate FSC=0.5 and FSC=0.143. **f**. 3D FSC plot and directional resolution histogram.

**Extended Data Figure 8.**
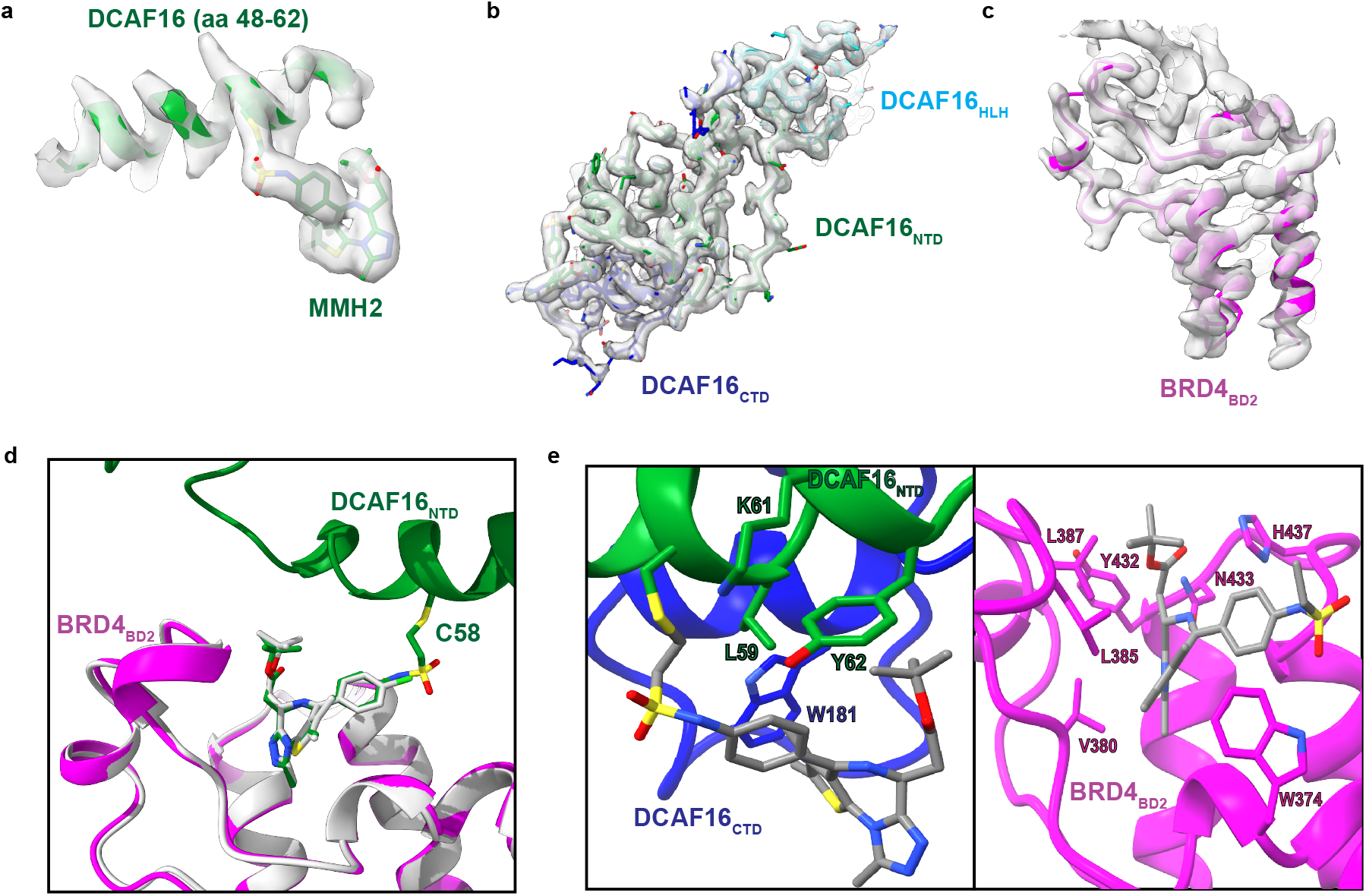
Map quality of the DCAF16-BRD4_BD2_-MMH2 interface. **a**. Cryo-EM density for DCAF16 containing _Cys58_ covalently bound to MMH2. Map contoured at 0.251. **b**.Cryo-EM density for DCAF16. **c**. Cryo-EM density for BRD4_BD2_. **d**. Overlay of MMH2 with JQ1 (PDB: 3ONI, in white). **e**. Key residues on DCAF16 (in green and blue) and BRD4_BD2_ (in magenta) close to MMH2.

**Extended Data Figure 9.**
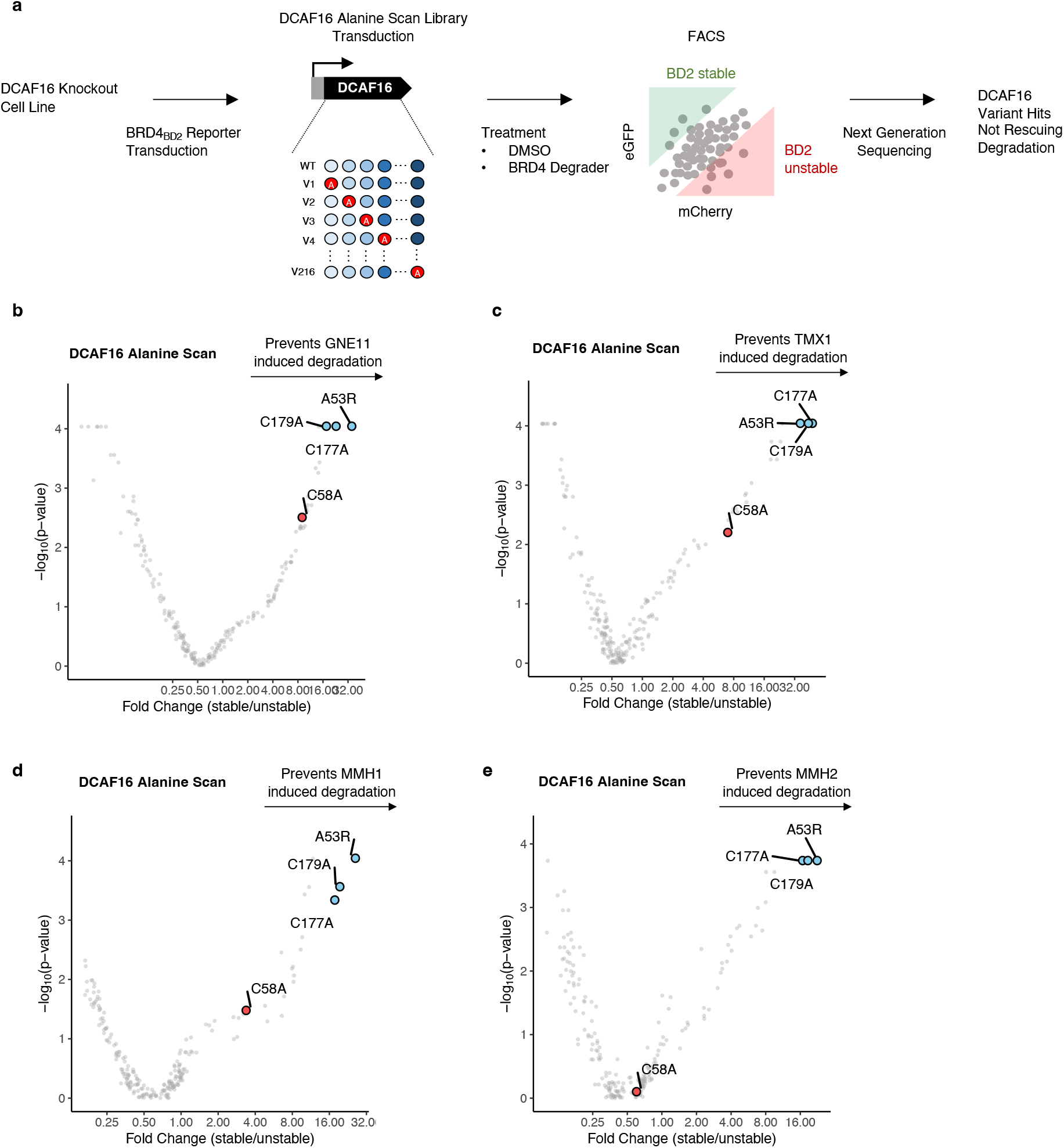
DCAF16 alanine-scanning reporter screen for BRD4 molecular glue degraders. **a**. Schematic of DCAF16 alanine mutagenesis screen for BRD4_BD2_-eGFP degradation in DCAF16 knockout K562 cells. **b**. DCAF16 alanine mutagenesis screen for BRD4_BD2_-eGFP stability in K562 cells treated with GNE11 at 1 μM for 16 h (n=3). **c**. DCAF16 alanine mutagenesis screen for BRD4_BD2_-eGFP stability in K562 cells treated with TMX1 at 1 μM for 16 h (n=3). **d**. DCAF16 alanine mutagenesis screen for BRD4_BD2_-eGFP stability in K562 cells treated with MMH1 at 0.1 μM for 16 h (n=3). **e**. DCAF16 alanine mutagenesis screen for BRD4_BD2_-eGFP stability in K562 cells treated with MMH2 at 0.1 μM for 16 h (n=3).

**Extended Data Figure 10.**
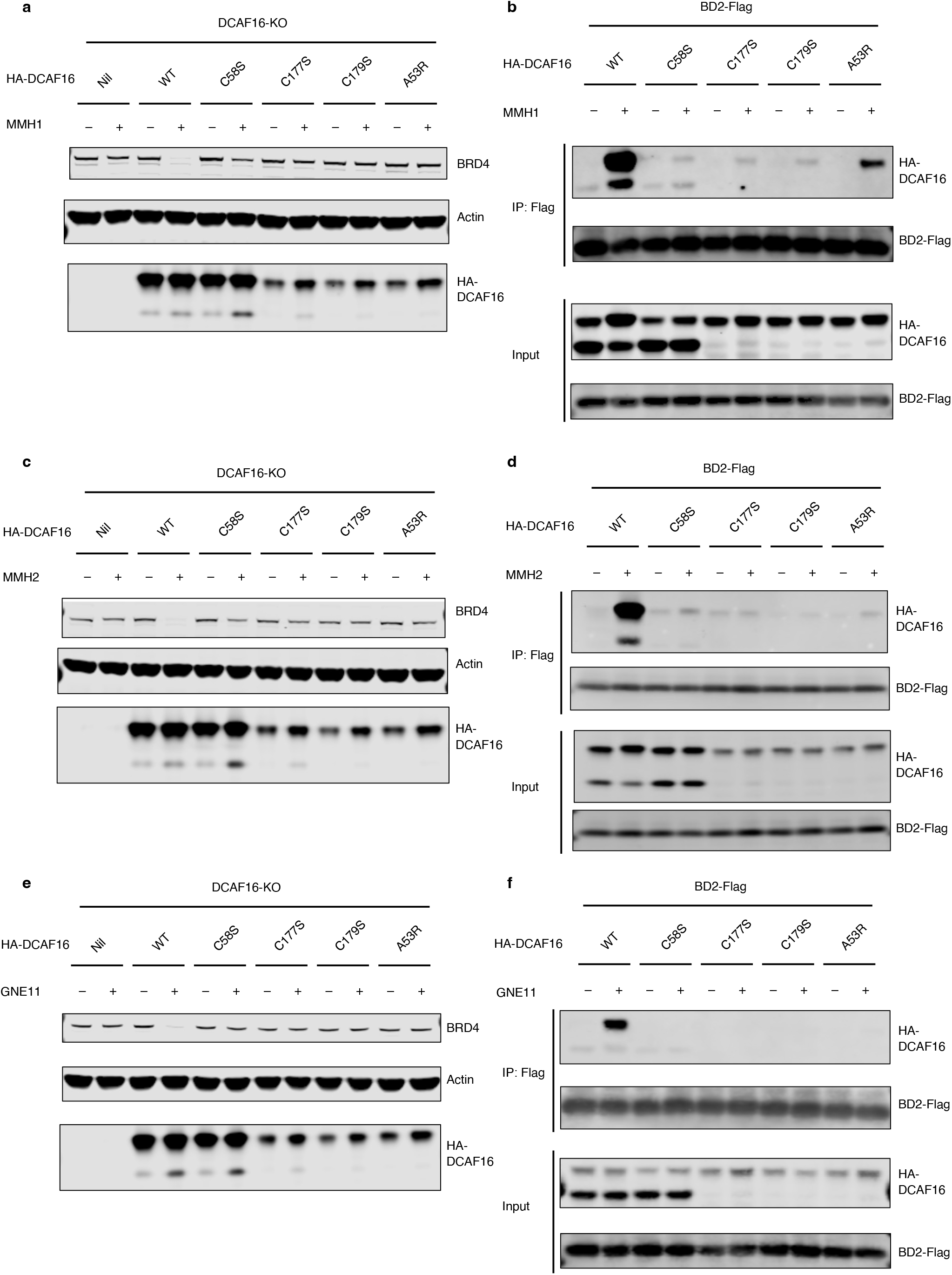
Cellular validation of representative DCAF16 mutants. **a**. Western blots of BRD4 degradation in DCAF16 knockout K562 cells that were transduced with indicated HA-DCAF16 mutants and treated with DMSO or MMH1 at 0.1 μM for 16 h. **b**. Flag immunoprecipitation (IP) followed by Western blots in the presence of DMSO or MMH1 at 0.1 μM from 293T cells transfected with indicated HA-DCAF16 mutants and BRD4_BD2_-Flag constructs. **c**. Western blots of BRD4 degradation in DCAF16 knockout K562 cells that were transduced with indicated HA-DCAF16 mutants and treated with DMSO or MMH2 at 0.1 μM for 16 h. **d**. Flag immunoprecipitation (IP) followed by Western blots in the presence of DMSO or MMH2 at 0.1 μM from 293T cells transfected with indicated HA-DCAF16 mutants and BRD4_BD2_-Flag constructs. **e**. Western blots of BRD4 degradation in DCAF16 knockout K562 cells that were transduced with indicated HA-DCAF16 mutants and treated with DMSO or GNE11 at 1 μM for 16 h. **f**. Flag immunoprecipitation (IP) followed by Western blots in the presence of DMSO or GNE11 at 1 μM from 293T cells transfected with indicated HA-DCAF16 mutants and BRD4_BD2_-Flag constructs.

**Extended Data Figure 11.**
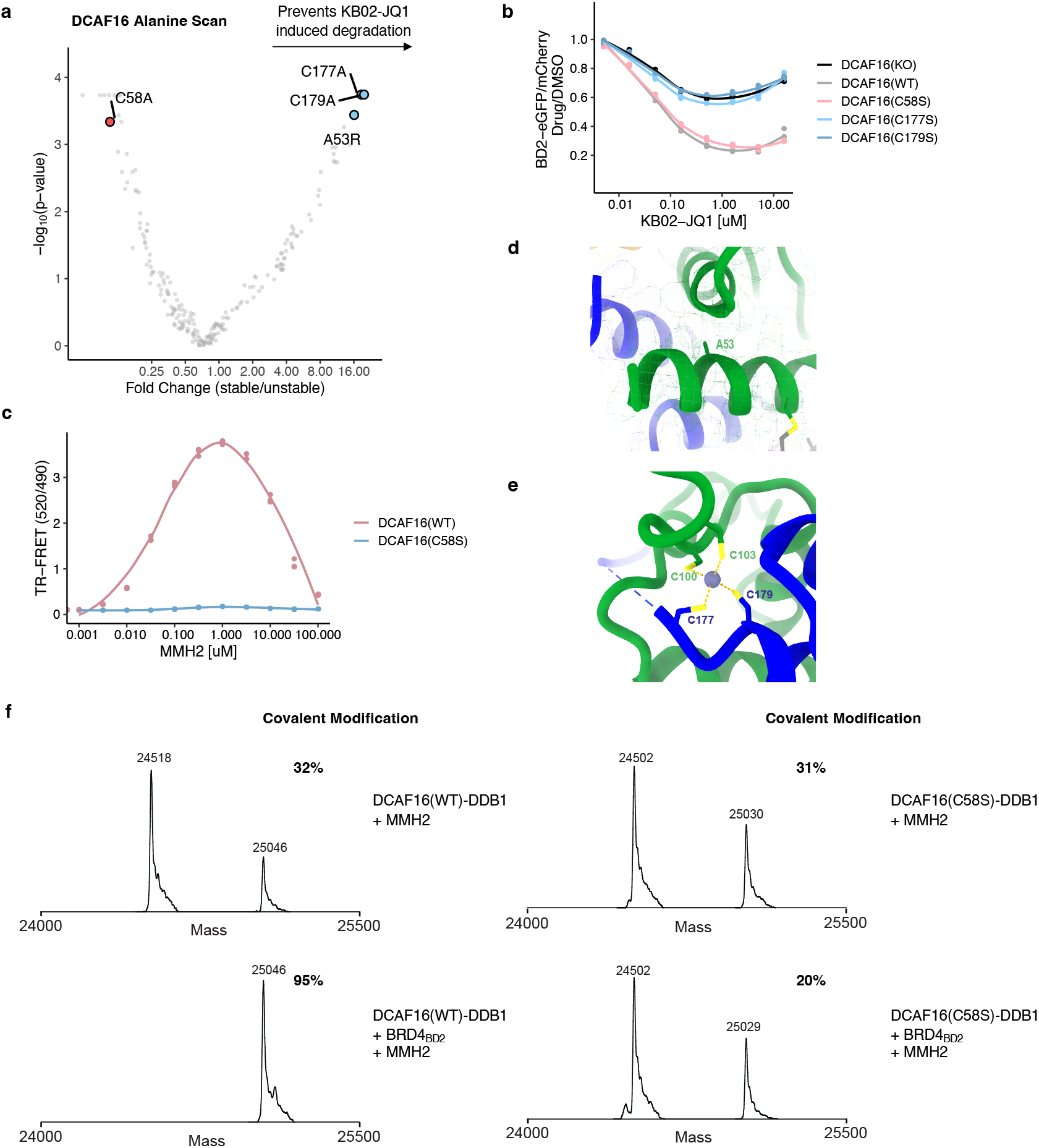
DCAF16 alanine-scanning reporter screen and validation. **a**. DCAF16 alanine mutagenesis screen for BRD4_BD2_-eGFP stability in K562 cells treated with KB02-JQ1 at 10 μM for 16 h (n=3). **b**. Flow analysis of BRD4_BD2_-eGFP degradation in DCAF16 knockout K562 cells transduced with indicated HA-DCAF16 mutants and treated with increasing concentrations of KB02-JQ1 for 16 h (n=3). **c**. TR-FRET signal for DDB1-DCAF16(WT)- or DDB1-DCAF16(C58S)-BODIPY to BRD4_BD2_-terbium with increasing concentrations of MMH2 (n=3). **d**. Close up of DCAF16 Ala53 orienting towards the hydrophobic core. **e**. Close up of DCAF16 Cys177 and Cys179 coordinating a structural zinc ion. **f**. Intact protein mass spectra of DDB1-DCAF16(WT) co-incubated with MMH2, DDB1-DCAF16(WT) co-incubated with MMH2 and BRD4_BD2_, DDB1-DCAF16(C58S) co-incubated with MMH2, or DDB1-DCAF16(C58S) co-incubated with MMH2 and BRD4_BD2_ at 4°C for 16 h.

**Extended Data Figure 12.**
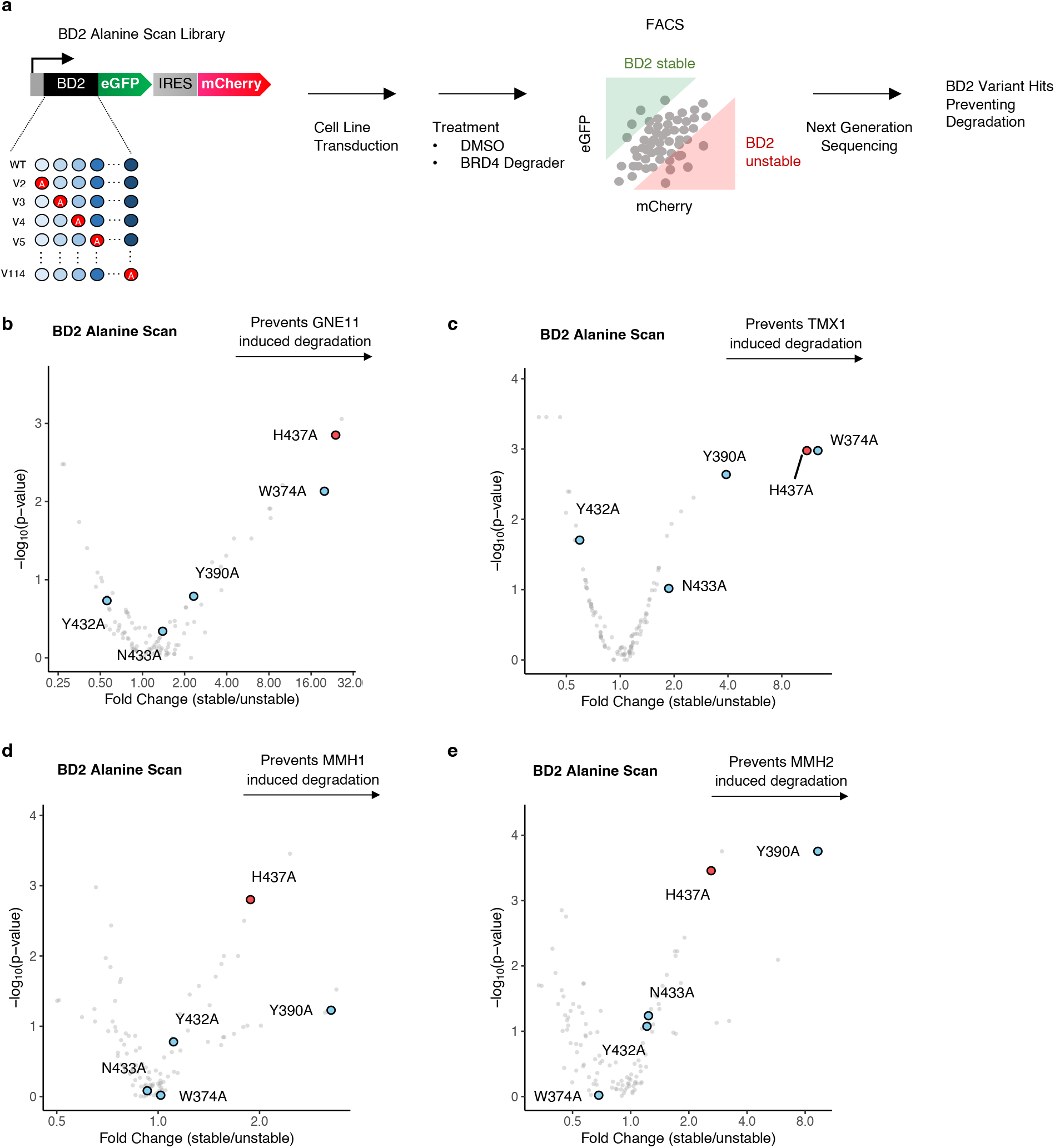
BRD4_BD2_ alanine-scanning reporter screen for BRD4 molecular glue degraders. **a**. Schematic of alanine mutagenesis degradation screen of the BRD4 _BD2_ domain in K562 cells. **b**. BD2 alanine mutagenesis screen for BRD4_BD2_-eGFP stability in K562 cells treated with GNE11 at 1 μM for 16 h (n=2). **c**. _BD2_ alanine mutagenesis screen for BRD4_BD2_-eGFP stability in K562 cells treated with TMX1 at 1 μM for 16 h (n=2). **d**. BD2 alanine mutagenesis screen for BRD4_BD2_-eGFP stability in K562 cells treated with MMH1 at 0.1 μM for 16 h (n=3). **e**. BD2 alanine mutagenesis screen for BRD4_BD2_-eGFP stability in K562 cells treated with MMH2 at 0.1 μM for 16 h (n=3).

**Extended Data Figure 13.**
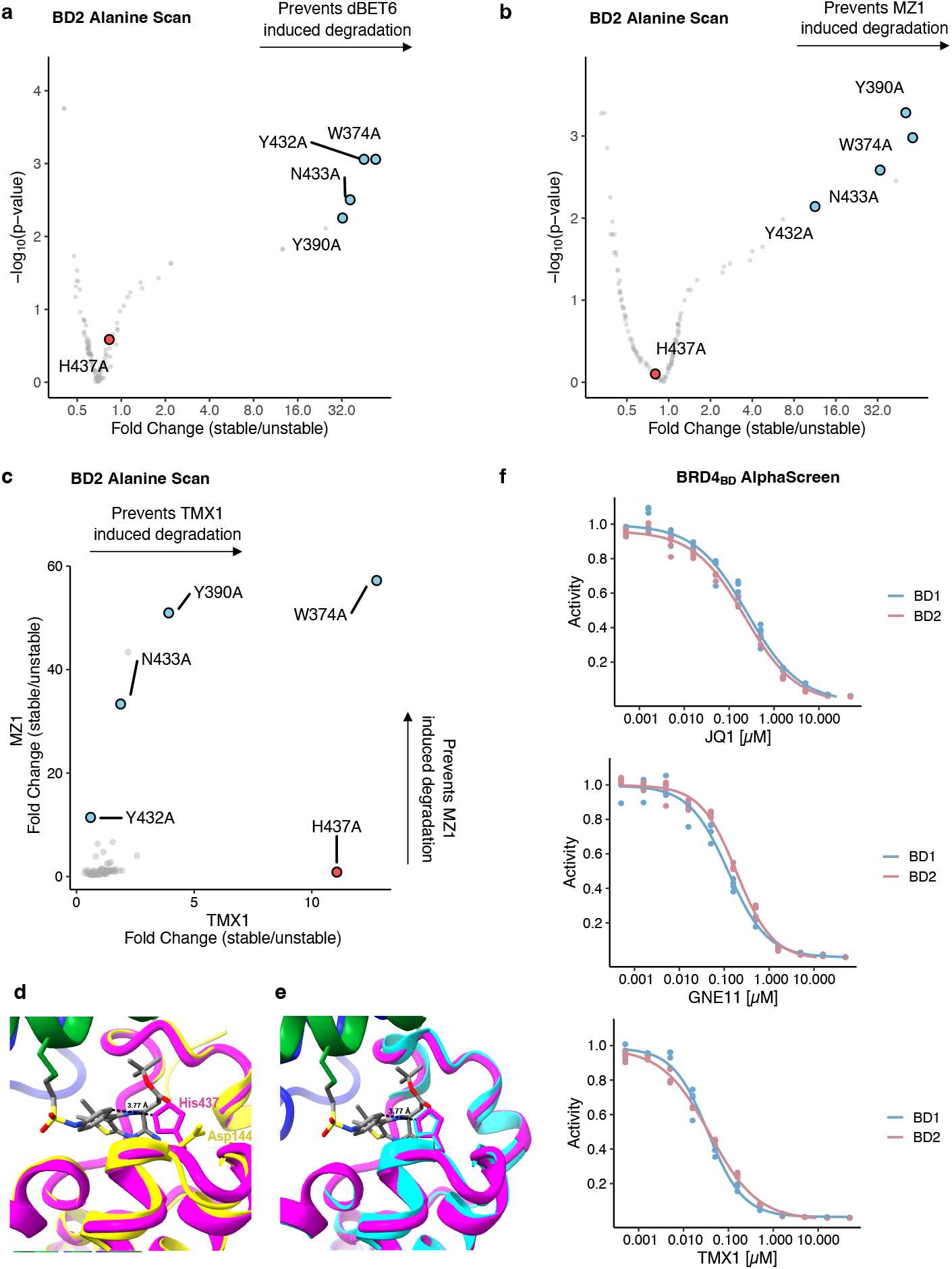
BRD4_BD2_ alanine-scanning reporter screen and mechanism of bromodomain selectivity. **a**. BD2 alanine mutagenesis screen for BRD4_BD2_-eGFP stability in K562 cells treated with dBET6 at 1 μM for 16 h (n=2). **b**. _BD2_ alanine mutagenesis screen for BRD4_BD2_-eGFP stability in K562 cells treated with MZ1 at 1 μM for 16 h (n=2). **c**. Correlation of fold change for two BRD4_BD2_ alanine mutagenesis screens. The *x* axis is a degradation screen for BRD4_BD2_-eGFP in K562 cells upon treatment with TMX1 at 1 μM for 16 h (n=2), and the *y* axis is another degradation screen for BRD4_BD2_-eGFP in K562 cells upon treatment with MZ1 at 1 μM for 16 h (n=2). **d**. Overlay of BRD4_BD1_ (PDB: 3MXF, in yellow) with BRD4_BD2_ (in magenta) showing a close-up of residues His437. When substituted for Asp144 in BRD4_BD1_, there is repulsion between Asp144 and the JQ1 carbonyl. **e**. Overlay of BRD2_BD2_ (PDB: 3ONI, in cyan) with BRD4_BD2_ (in magenta) showing a close-up of residues His437 and the corresponding His433 in BRD2_BD2_. **f**. AlphaScreen competitive assay of JQ1, GNE11, and TMX1 to quantify the drug’s inhibition of binding between biotinylated-JQ1 and His-tagged BRD4_BD1_ or BRD4_BD2_ (n=4).

**Extended Data Table 1.**
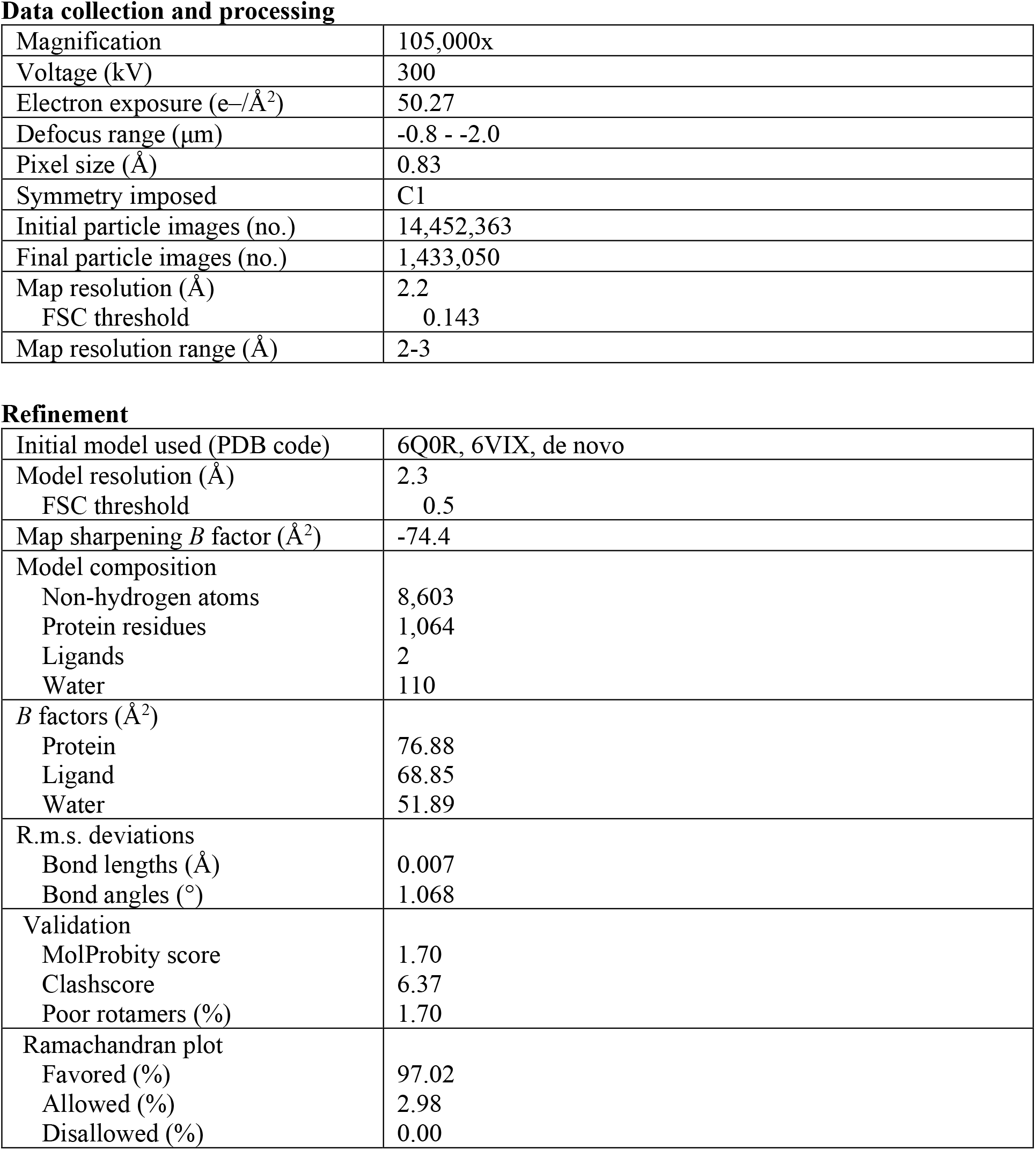
Cryo-EM data collection, refinement, and validation statistics. DCAF16-DDB1ΔB-DDA1-MMH2-BRD4_BD2_ (EMDB-29714) (PDB 8G46)

Data collection and processing
|  |  |
| --- | --- |
| Magnification | 105,000x |
| Voltage (kV) | 300 |
| Electron exposure (e-/Å<sup>2</sup>) | 50.27 |
| Defocus range (μm) | -0.8 - -2.0 |
| Pixel size (Å) | 0.83 |
| Symmetry imposed | C1 |
| Initial particle images (no.) | 14,452,363 |
| Final particle images (no.) | 1,433,050 |
| Map resolution (Å) | 2.2 |
| FSC threshold | 0.143 |
| Map resolution range (Å) | 2-3 |
| Refinement |  |
| Initial model used (PDB code) | 6Q0R, 6VIX, de novo |
| Model resolution (Å) | 2.3 |
| FSC threshold | 0.5 |
| Map sharpening *B* factor (Å<sup>2</sup>) | -74.4 |
| Model composition |  |
| Non-hydrogen atoms | 8,603 |
| Protein residues | 1,064 |
| Ligands | 2 |
| Water | 110 |
| *B* factors (Å<sup>2</sup>) |  |
| Protein | 76.88 |
| Ligand | 68.85 |
| Water | 51.89 |
| R.m.s. deviations |  |
| Bond lengths (Å) | 0.007 |
| Bond angles (°) | 1.068 |
| Validation |  |
| MolProbity score | 1.70 |
| Clashscore | 6.37 |
| Poor rotamers (%) | 1.70 |
| Ramachandran plot |  |
| Favored (%) | 97.02 |
| Allowed (%) | 2.98 |
| Disallowed (%) | 0.00 |

